# Trans-generational inheritance of centromere identity requires the CENP-A N-terminal tail in the *C. elegans* maternal germ line

**DOI:** 10.1101/2020.10.05.325985

**Authors:** Reinier F. Prosée, Joanna M. Wenda, Caroline Gabus, Kamila Delaney, Francoise Schwager, Monica Gotta, Florian A. Steiner

**Affiliations:** Department of Molecular Biology and Institute for Genetics and Genomics in Geneva, Section of Biology, Faculty of Sciences, University of Geneva, 1211 Geneva, Switzerland; Department of Cell Physiology and Metabolism and Institute for Genetics and Genomics in Geneva, Faculty of Medicine, University of Geneva, 1211 Geneva, Switzerland

**Keywords:** Centromere, CENP-A, C. elegans, KNL-2, trans-generational inheritance

## Abstract

Centromere protein A (CENP-A) is a histone H3 variant that defines centromeric chromatin and is essential for centromere function. In most eukaryotes CENP-A-containing chromatin is epigenetically maintained, and centromere identity is inherited from one cell cycle to the next. In the germ line of the holocentric nematode *Caenorhabditis elegans*, this inheritance cycle is disrupted. CENP-A is removed at the mitosis-to-meiosis transition and is established *de novo* on chromatin during diplotene of meiosis I. Here we show that the N-terminal tail of CENP-A is required for the *de novo* establishment of centromeres, but dispensable for centromere maintenance during embryogenesis. Worms homozygous for a CENP-A tail deletion maintain a functional centromere during development, but give rise to inviable offspring because they fail to re-establish centromeres in the maternal germ line. We identify the N-terminal tail of CENP-A as a critical domain for the interaction with the conserved kinetochore protein KNL-2, and argue that this interaction plays an important role in setting centromere identity in the germ line. We conclude that centromere establishment and maintenance are functionally distinct in *C. elegans*.

## Introduction

Centromeres are crucial for the segregation of chromosomes into the daughter cells during cell division. They hold the sister chromatids together after DNA replication and serve as the sites of kinetochore formation. During mitosis, the kinetochore attracts and binds microtubules that pull the sister chromatids towards opposing spindle poles. In most species, the centromere is defined by the histone variant CENP-A that forms the chromatin foundation for building the kinetochore and acts as an epigenetic mark to maintain centromere identity [1,2]. CENP-A is almost universally conserved, but differs in amino acid sequence between different taxa, especially in the N-terminal tail [3–6]. The C-terminal histone fold domain (HFD) is more conserved, since it incorporates into the histone octamer and is therefore under structural constraints [7].

The incorporation and maintenance of CENP-A at the centromere is a highly regulated process [8,9]. During DNA replication, CENP-A levels are diluted by half and have to be replenished with newly synthesized CENP-A [10]. Most studies investigating CENP-A loading have been performed in cultured mitotic cells, where it is possible to tightly regulate the cell cycle. Although the exact mechanisms differ slightly between different species, the basic components for CENP-A loading and maintenance in mitosis are well conserved [2]. CENP-C and the MIS18 complex recognize CENP-A nucleosomes and are required for priming centromeric sites for new CENP-A deposition [11–14]. CENP-A-specific histone chaperones (HJURP/CAL1/Scm3) then incorporate new CENP-A at centromeres, thereby maintaining CENP-A on chromatin [15–17]. Placeholder nucleosomes, containing H3 or the histone variant H3.3, can mark the location where new CENP-A will be loaded [3,18], and active transcription displaces these nucleosomes to allow for the deposition of the CENP-A nucleosomes [19–21].

How centromeres are inherited from one generation to the next in multicellular organisms is less well understood [22]. However, the inheritance of CENP-A protein into the zygote is crucial for centromere functioning in the next generation prior to the onset of embryonic transcription. This inheritance is achieved either through stability of the CENP-A nucleosomes or active recycling of CENP-A during meiosis [23–25]. In organisms that inherit CENP-A protein only on the paternal (e.g. *Arabidopsis thaliana*) or the maternal (e.g. *C. elegans*) chromosomes, the ‘template-governed’ model of centromere inheritance has been disputed, because the paternal or maternal chromosomes devoid of CENP-A need to establish centromeres *de novo* after fertilization and prior to the first embryonic cell division [26–28]. An additional interruption of the templated inheritance of CENP-A occurs in the adult hermaphrodite germ line of *C. elegans*, where CENP-A is removed from chromatin at the mitosis-to-meiosis transition and reloaded during diplotene of meiosis I [28]. How this *de novo* establishment of CENP-A is regulated remains unclear.

The functional *C. elegans* CENP-A homologue is called HCP-3, but in this study we will refer to it as CENP-A for clarity. *C. elegans* chromosomes are holocentric, with CENP-A distributed along the length of the entire chromosomes [29]. CENP-A is essential for mitotic cell divisions in the distal germ line and during development, but dispensable for chromosome segregation during meiosis [28]. Chromatin association of CENP-A during mitosis depends on the MIS18BP1 homologue KNL-2 [30] and on the general histone chaperone LIN-53, a homologue of RbAp46/48 [31].

In this study, we demonstrate that the N-terminus of CENP-A mediates interaction with KNL-2 *in vitro* and *in vivo*. Surprisingly, removing the N-terminal tail of CENP-A does not affect centromere function during embryogenesis, and the kinetochore-null phenotype associated with loss of centromeres only appears at the first embryonic division of the next generation. We find that correctly established centromere identity, rather than the presence of the full-length CENP-A protein, is essential for mitosis during development. Our results suggest that in *C. elegans* the CENP-A N-terminal tail has acquired an important function in re-establishing the centromere in each generation in the adult germ line, but is dispensable for centromere maintenance.

## Results

### The CENP-A N-terminal tail interacts with the central domain of KNL-2

In *C. elegans*, CENP-A is chromatin-associated in dividing cells throughout development, but is removed and re-established, respectively, during different stages of the adult hermaphrodite germ line [28] (Fig 1). In embryos, KNL-2 and CENP-A co-localize and are interdependent for chromatin association, and depletion of either KNL-2 or CENP-A leads to a kinetochore-null phenotype [30,32]. CENP-A and KNL-2 also show overlapping germ line distributions (Fig 1). We therefore hypothesized that KNL-2 may play a role in the *de novo* establishment of centromeres in the germ line.

**Fig 1.**
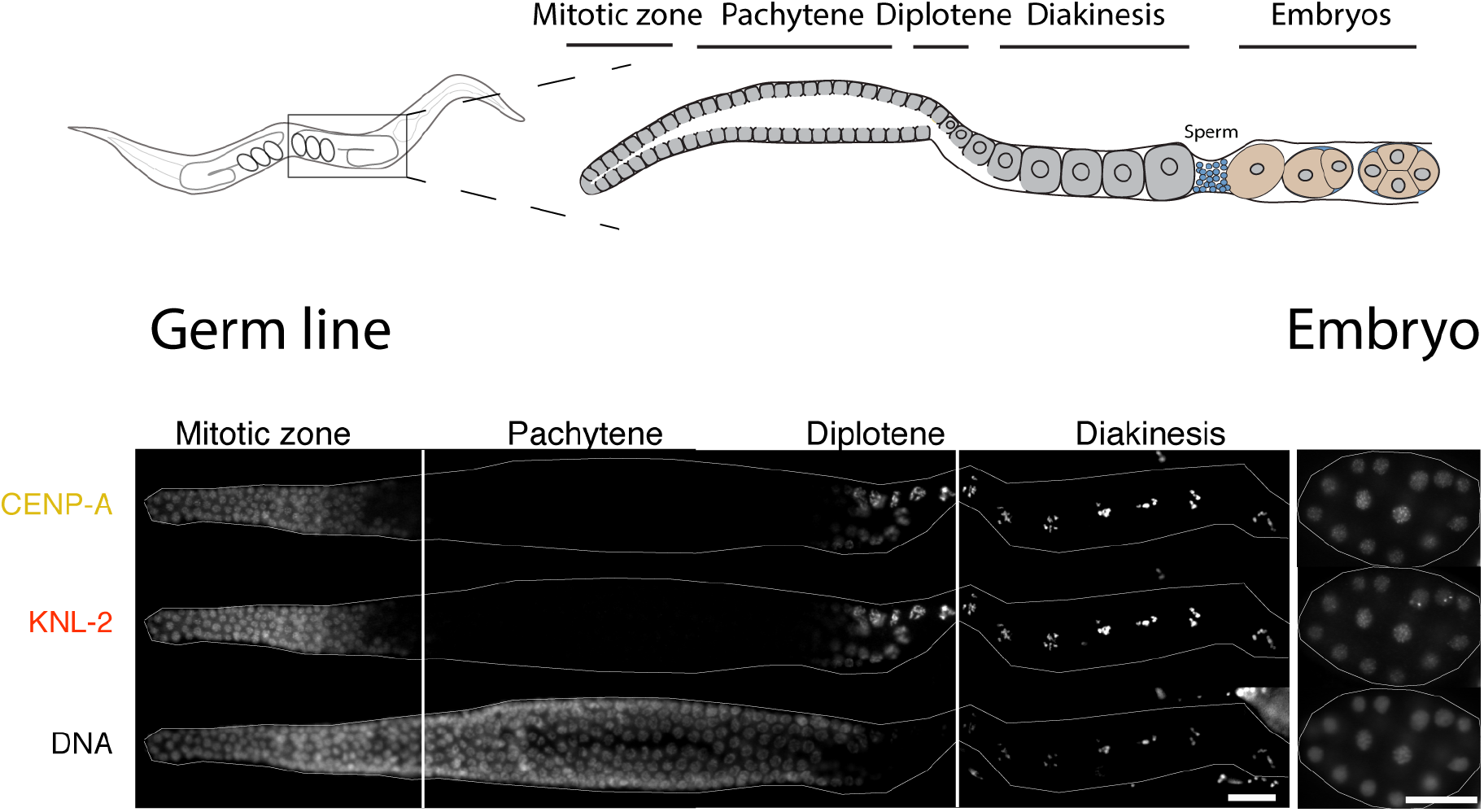
Dynamics of KNL-2 and CENP-A in the *C. elegans* germ line. Top, cartoon images of an adult hermaphrodite *C. elegans* and the different stages of the germ line. Bottom, immunofluorescence (IF) images of CENP-A and KNL-2 in adult hermaphrodite germ lines and embryos, showing the removal at the mitosis-meiosis transition and the reappearance at the diplotene stage. Scale bars represent 20 μm.

We first investigated the interplay between CENP-A and KNL-2. We used a yeast two-hybrid (Y2H) assay to determine the minimal regions of both proteins that are required for their interaction. The interaction between full-length CENP-A and KNL-2 is barely detectable (Fig 2A, left), possibly due to improper folding of the proteins. Next, we tested the CENP-A histone fold domain (HFD) or the N-terminal tail in combination with full-length KNL-2. While we observed no interaction between the CENP-A HFD and KNL-2, the interaction between the CENP-A N-terminal tail and KNL-2 supported yeast growth on selective medium (SD-LTH containing 5mM 3AT; Fig 2A, left). We then tested constructs covering the N-terminal (aa 1-268), central (aa 269-470) and C-terminal (aa 471-877) regions of KNL-2. We only observed yeast growth on selective medium for the central region (aa 269-470), as was reported in an earlier study [33] (Fig 2A, middle). This part of KNL-2 does not contain any annotated domains and is predicted to be mostly unstructured. We next tested different parts of the CENP-A N-terminal tail for interaction with the KNL-2 central region, and found that aa 1-122 are sufficient to promote yeast growth on selective medium (Fig 2A, right). We confirmed the interaction of the CENP-A N-terminal tail with the KNL-2 central region *in vitro*, using recombinant proteins (S1 Fig).

**Fig 2.**
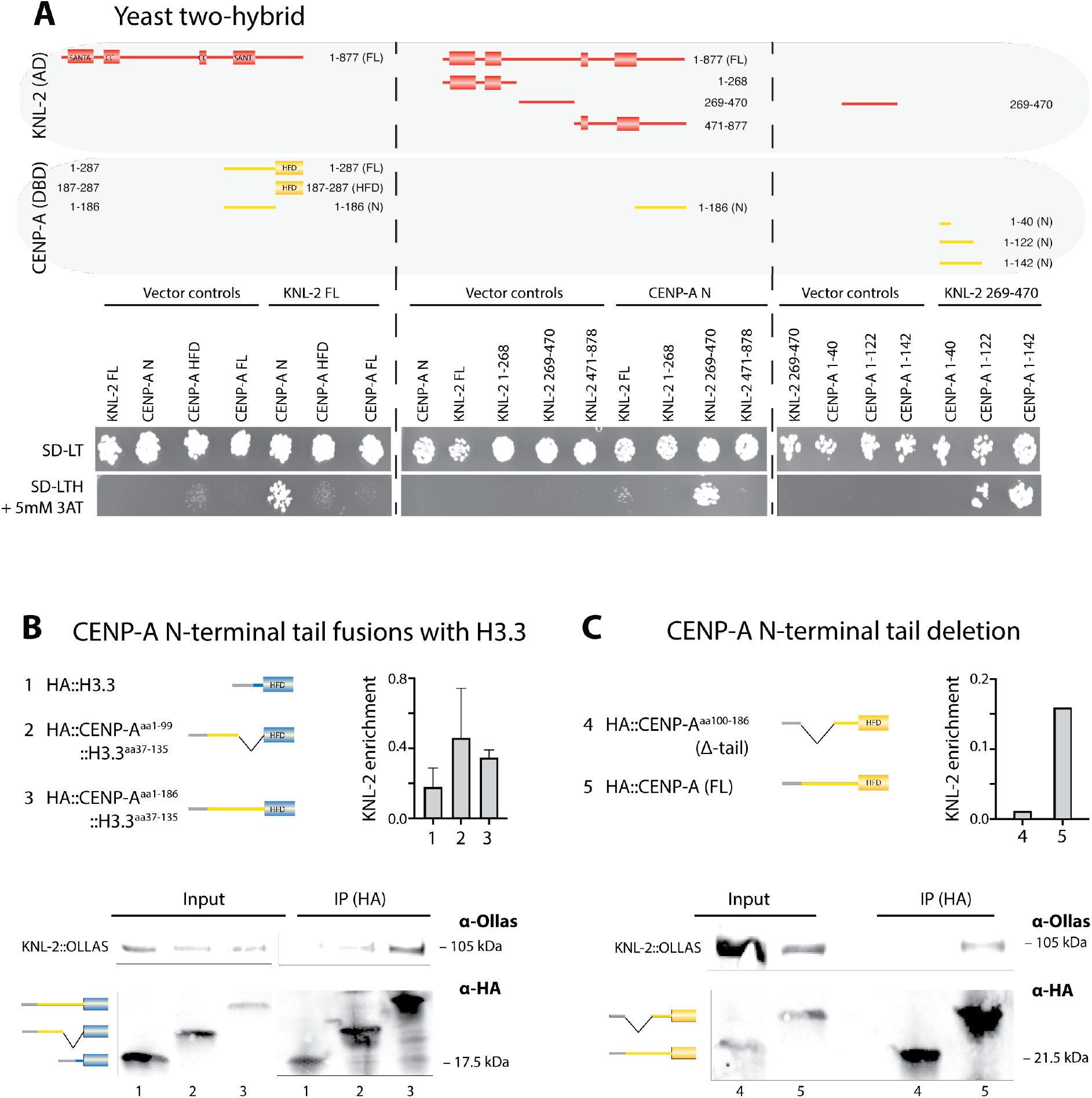
The CENP-A N-terminal tail interacts with the KNL-2 central domain. (A) Yeast two-hybrid analysis of the interaction between CENP-A and KNL-2. Cartoons of the different CENP-A and KNL-2 fragments used (top), showing the annotated SANTA, SANT/Myb and coiled-coil (CC) domains in KNL-2 and histone-fold domain (HFD) in CENP-A. The interaction between the constructs were determined by assaying growth of yeast cells on selective medium (SD-LTH, containing 5mM 3AT). Vector-only constructs and non-selective plates (SD-LT) were used as negative and positive controls for growth, respectively. Left, full-length KNL-2 interaction with different parts of CENP-A. Center, interaction of the CENP-A N-terminal tail with different regions of KNL-2. Right, interaction of different CENP-A N-terminal tail fragments with the KNL-2 central region. (B) Co-IP experiments with KNL-2 and CENP-A::H3.3 chimeras. In cartoons, CENP-A is shown in yellow, H3.3 in blue and HA in gray. Representative western blots showing IPs of HA-tagged H3.3, CENP-A^aa1-99^::H3.3^aa37-135^, and CENP-A^aa1-186^::H3.3^aa37-135^ from embryonic extracts. HA-tagged constructs were IPed and detected using an anti-HA antibody. OLLAS-tagged KNL-2 was detected using an anti-OLLAS antibody in input and IP samples. The bar graph shows a quantification of relative KNL-2 enrichment in two independent experiments. Error bars represent the standard deviation of the mean. (C) Co-IP experiments with KNL-2 and CENP-A N-terminal tail truncation. In cartoons, CENP-A is shown in yellow and HA in gray. Representative western blot showing IPs of HA-tagged CENP-A (FL), or CENP-A^aa100-287^ (Δ-tail) from embryonic extracts. HA-tagged constructs were IPed and detected using an anti-HA antibody. OLLAS-tagged KNL-2 was detected using an anti-OLLAS antibody in input and IP samples. The bar graph shows a quantification of relative KNL-2 enrichment in the western blot shown below. KNL-2 enrichment was measured by determining the input-corrected ratio of KNL-2 IP-signal to CENP-A IP-signal in (B) and (C).

Next, we tested the interaction of the CENP-A N-terminal tail with KNL-2 *in vivo*. In order to target different parts of the CENP-A N-terminal tail to chromatin, we fused them to the HFD of HIS-72, a non-essential, ubiquitously expressed histone H3.3 homologue [34,35]. We tested the interaction of these constructs with KNL-2 by pull-down experiments of HA-tagged fusion proteins from embryonic extracts. We found that replacing the N-terminal tail of HIS-72 (aa 1-36) with the entire CENP-A N-terminal tail (aa 1-186) or with the first 99 aa of the CENP-A N-terminal tail was sufficient to mediate interaction with KNL-2 in co-IP experiments, while HIS-72 interacted with KNL-2 two-fold less efficiently than the CENP-A N-terminal tail-H3.3 chimeras (Fig 2B).

We then removed the part of the endogenous *hcp-3* gene encoding the first 99 amino acids of the CENP-A N-terminal tail using the CRISPR/Cas9 system (CENP-A^aa100-287^; called CENP-A Δ-tail for simplicity). We maintained this truncation allele heterozygous over the wild type allele in the context of a genetic balancer. In this balancer strain, the balanced allele with the full-length copy of *hcp-3* (CENP-A FL) is linked to a transgene expressing pharyngeal GFP, thus visually allowing us to distinguish worms heterozygous and homozygous for the CENP-A N-terminal tail deletion. *C. elegans* populations mostly consist of hermaphrodites that produce both oocytes and sperm and self-fertilize to give rise to the next generation, although males that only produce sperm and rely on cross-fertilizing hermaphrodites to give rise to offspring also exist. In this study, we use the heterozygous (CENP-A FL/Δ-tail) worms as parental generation (P0), and label the first and second subsequent offspring generations as F1 and F2, respectively. F1 homozygotes for the full-length copy of CENP-A are embryonic lethal due to aneuploidy of the balanced chromosomes. We used this balancer strain to investigate the interaction of KNL-2 with full-length and truncated CENP-A *in vivo*. IP of the CENP-A Δ-tail protein did not co-precipitate KNL-2, while KNL-2 was easily detectable in IPs of full-length CENP-A (Fig 2C).

We conclude that the CENP-A N-terminal tail is both necessary and sufficient for the interaction with KNL-2. Based on our Y2H assay and the *in vitro* experiments, the central region of KNL-2 mediates this interaction.

### The N-terminal tail of CENP-A is dispensable during development

CENP-A and KNL-2 are both essential proteins, and loss-of-function mutations or depletion by RNAi results in early embryonic lethality due to chromosome segregation errors [30,36]. Based on the previously reported interdependence of CENP-A and KNL-2 in maintaining the centromere, we expected to observe an embryonic lethal phenotype when interrupting their interaction. However, we found that in the F1 generation, hermaphrodites homozygous for CENP-A Δ-tail were fully viable (Fig 3A). These worms looked and developed indistinguishably from wild type worms and showed no obvious germline proliferation defects. In addition, homozygous F1 CENP-A Δ-tail/Δ-tail males were viable and fertile (S2A Fig). However, the F2 embryos laid by the homozygous F1 hermaphrodites were all inviable due to severe chromosome segregation defects (Fig 3A, B). Instead of the expected F1 embryonic lethal (*emb*) phenotype, the CENP-A N-terminal tail deletion thus causes a maternal effect lethal (*mel*) phenotype.

**Fig 3.**
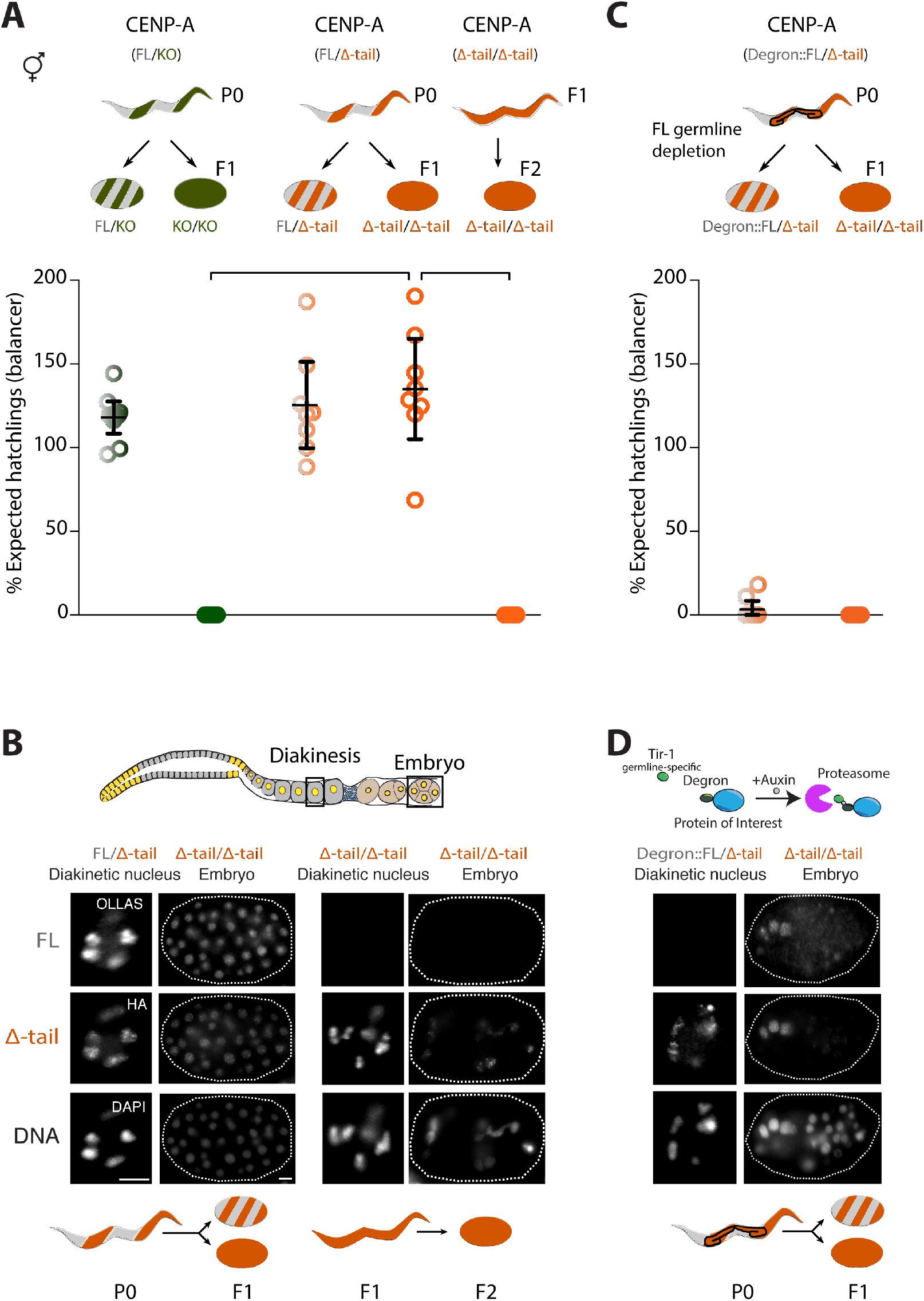
The CENP-A N-terminal tail is dispensable for mitosis during development but required in the germ line. (A, B) Phenotypic consequences of the CENP-A N-terminal tail deletion in presence or absence of CENP-A full-length protein in the maternal germ line. (A) Quantification of viable offspring of worms carrying CENP-A full-length (FL, gray), CENP-A null mutation (KO, green) or CENP-A tail truncation (Δ-tail, orange) alleles. A balancer allele (hT2) was used to maintain heterozygous strains. In these balancer strains, CENP-A full-length homozygotes are embryonic lethal due to the balancer alleles and are therefore not shown. Top, cartoon of maternal heterozygous (striped) or homozygous (full) genotypes as well as genotypes of the offspring. Bottom, quantification of the viable offspring as a percentage of the expected hatchlings in the context of the balancer allele. Error bars show the 95% confidence interval of the mean. Connecting lines highlight the most relevant comparisons between the different genotypes. (B) Cartoon of an adult germ line, with imaged regions highlighted with black boxes. IF images of both the CENP-A FL and CENP-A Δ-tail protein in the maternal germ line (diakinesis) and embryos. Left, maternal FL/Δ-tail P0 giving rise to either FL/Δ-tail or Δ-tail/Δ-tail F1 offspring (indistinguishable as embryos). Right, maternal Δ-tail/Δ-tail F1 giving rise to Δ-tail/Δ-tail F2 offspring. Cartoon images of adult worms and embryos with the different genotypes as described in (A). (C, D) Depletion of the CENP-A full-length protein in the germ line of FL/Δ-tail heterozygous worms (P0), using the AID system. (C) Top, cartoon of maternal genotype and genotypes of the offspring. Bottom, quantification of the viable offspring as a percentage of the expected hatchlings in the context of the balanced allele. Color codes as in (A). (D) Cartoon of the auxin-induced degron (AID) technique. IF images of both the CENP-A FL and CENP-A Δ-tail protein in the maternal germ line (diakinesis) and embryos. Maternal FL/Δ-tail P0 upon AID-mediated depletion of FL in the germ line, giving rise to either FL/Δ-tail or Δ-tail/Δ-tail F1 offspring (indistinguishable as embryos). Cartoon images of adult worms and embryos with the different genotypes as described in (A). Scale bars in (B, D) represent 5 μm.

In *C. elegans*, the CENP-A protein is exclusively inherited through the maternal germ line [26,28]. The *mel* phenotype caused by the CENP-A N-terminal tail deletion could potentially be explained by maternal contribution of the full-length protein from the P0 adults to the F1 embryos, as the maternal germ line (P0) is heterozygous for the truncation. This would require the maternal load to support the cell divisions during embryonic and larval stages as well as the germline proliferation in L4 and adult stages of the CENP-A Δ-tail F1 homozygotes. However, this is unlikely to be the case, since a strain containing a balanced *hcp-3* deletion produces homozygous F1 offspring with an embryonic lethal phenotype, even though these embryos presumably receive the same dose of full-length CENP-A from their maternal heterozygous germ line as those of the CENP-A Δ-tail strain (Fig 3A). Moreover, we did not detect any full-length CENP-A in the germ line of adult CENP-A Δ-tail F1 worms by immunostaining (Fig 3B).

To exclude the possibility that the truncated version of CENP-A stabilizes undetectable amounts of full-length CENP-A that could support development, we depleted full-length CENP-A during embryogenesis using the auxin-inducible degron (AID) system [37], with a TIR1 specific to the soma. Somatic depletion of full-length CENP-A leads to embryonic or larval lethality in wild type animals (S2B Fig). In contrast, homozygous CENP-A Δ-tail F1 embryos hatch and develop normally into adult worms, suggesting that there are no trace amounts of full-length CENP-A present during development, and that CENP-A Δ-tail protein is capable of supporting mitosis in the absence of full-length CENP-A (S2B Fig). Together, these results suggest that the CENP-A N-terminal tail is dispensable for mitosis during embryogenesis and germline proliferation.

### Full-length CENP-A is required for *de novo* centromere establishment in the maternal germ line

To explain the difference between the fully viable F1 CENP-A Δ-tail homozygotes and the embryonic lethal F2 CENP-A Δ-tail homozygotes, we considered how centromeres are established in the germ line and inherited into the embryos. The F1 animals inherit their centromeres from heterozygous P0 CENP-A Δ-tail germ lines, where full-length CENP-A is present during the centromere establishment step in prophase of meiosis I. The F2 animals, however, inherited their centromere from homozygous F1 CENP-A Δ-tail germ lines, where no full-length CENP-A is present. We therefore hypothesized that the CENP-A N-terminal tail is essential for centromere *de novo* establishment in the germ line. If this is the case, depletion of the full-length CENP-A from the germ line of heterozygous P0 CENP-A FL/Δ-tail animals should result in embryonic lethality of all F1 offspring, including the ones homozygous for the CENP-A Δ-tail. To test this, we depleted the full-length CENP-A protein in the heterozygous generation (P0) by using AID with a germline-specific TIR1 driver. We indeed observed severe chromosome missegregation in all F1 embryos, which mimics the *emb* phenotype observed in the F2 homozygous generation (Fig 3C, D).

Consistent with a role of full-length CENP-A in centromere establishment in the maternal germ line, crossing in a CENP-A full-length gene copy through the paternal germ line does not rescue F2 embryonic lethality (S2A Fig). This result supports the observation that there is no paternal contribution of CENP-A protein to the zygote [26], and shows that regaining a full-length CENP-A gene copy in the embryo is not sufficient for viability if no full-length protein was present in the maternal germ line. We conclude that the presence of full-length CENP-A in the maternal germ line is required for viability of the offspring.

### CENP-A Δ-tail F2 embryos show aberrant centromeres and a kinetochore-null phenotype

We next investigated the centromere formation defects in more detail. We decided to use the striking bi-orientation of CENP-A and KNL-2 on the metaphase plate in mitotic cells (both in the distal mitotic zone of the germ line and in embryos) as a visible read-out of a correctly assembled centromere. As expected, mitotic cells in the germ line of P0 heterozygotes (CENP-A FL/Δ-tail) show this bi-orientation (Fig 4A). In the F1 homozygote generation (CENP-A Δ-tail/Δ-tail) bi-orientation is visible in embryonic mitotic cells and in adult mitotic germ cells (Fig 4A). This observation supports the model that once centromere identity is established in the heterozygous germ line of the P0, it can be inherited into the adult germ line of the homozygous F1 (Fig 4A). In F2 embryonic mitotic cells, localization of both CENP-A and KNL-2 becomes disordered (Fig 4A). CENP-A still appears on chromatin, but is no longer bi-oriented. KNL-2 hardly remains present on chromatin and instead clusters in large dots that co-localize with the microtubules (Fig 4A, S3 Fig; S1 and S2 Videos).

**Fig 4.**
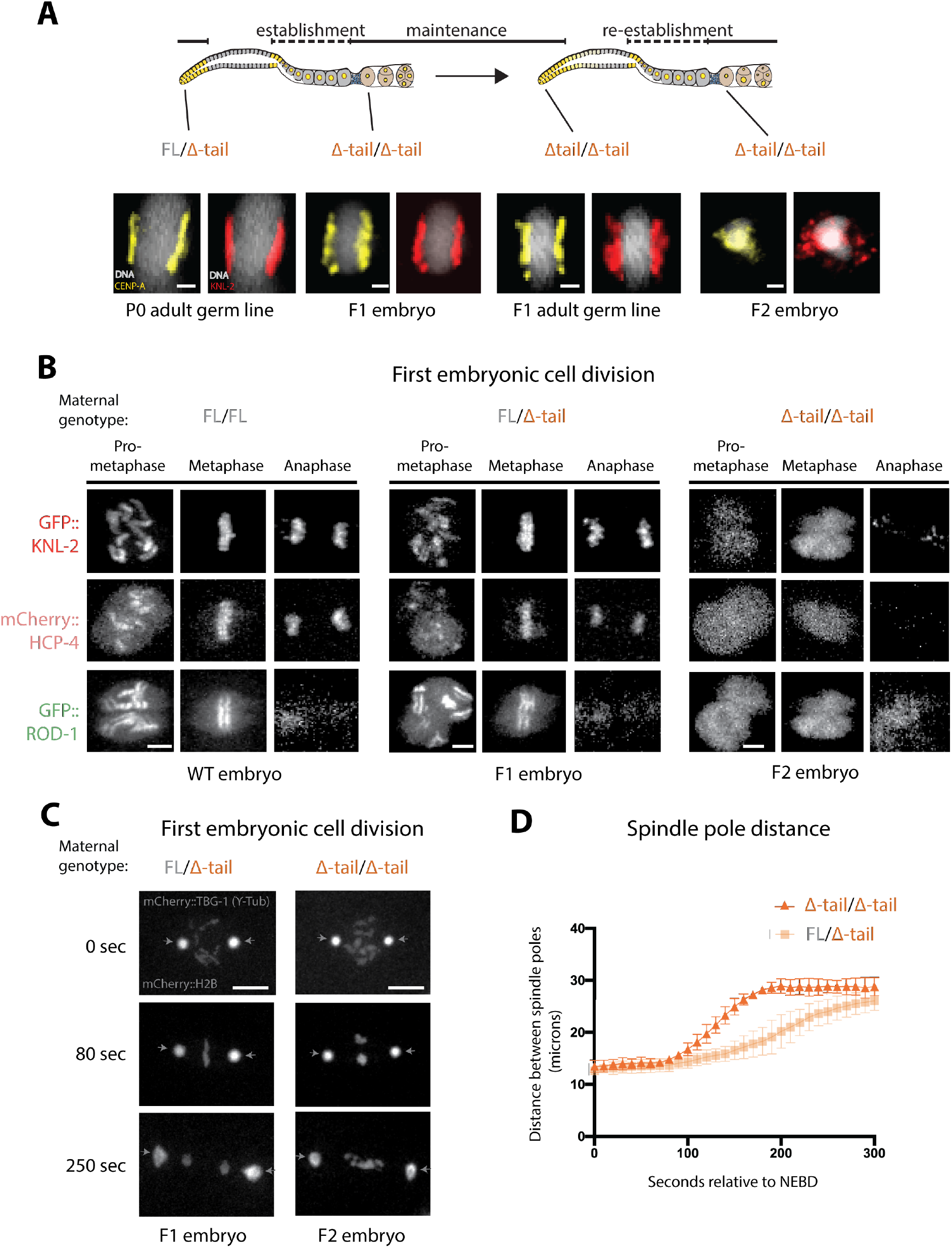
Centromere identity and function is maintained for one generation upon deletion of the CENP-A N-terminal tail. (A) Centromere bi-orientation in metaphase cells. IF images of KNL-2 and CENP-A Δ-tail in metaphase cells in the mitotic zone of the adult hermaphrodite germ line or in early embryos for the different generations (P0, F1, F2) of the CENP-A Δ-tail strain. (B) Still images for the indicated stages of mitosis of recordings in embryos derived from FL/FL, FL/Δ-tail or Δ-tail/Δ-tail maternal germ lines. KNL-2 (GFP), HCP-4 (mCherry), or ROD-1 (GFP) were fluorescently labeled. (C) Still images of recordings in embryos derived from FL/Δ-tail or Δ-tail/Δ-tail maternal germ lines, showing GFP-histone H2B (chromosomes) and GFP–γ-tubulin (spindle poles). Time is given in seconds relative to nuclear envelope breakdown. The arrows indicate the location of the spindle poles. (D) Quantification of spindle pole separation kinetics during the first embryonic cell division in embryos derived from FL/Δ-tail or Δ-tail/Δ-tail maternal germ lines. Error bars represent the standard deviation of the mean. Scale bars represent 1 μm (A), 4 μm (B) or 10 μm (C).

The mislocalization of both CENP-A and KNL-2 in the first embryonic cell division of F2 homozygote (CENP-A Δ-tail/Δ-tail) embryos leads to a dramatic missegregation of chromosomes, which explains the penetrant embryonic lethal phenotype (Figs 3A and 4A; S3 and S4 Videos). Live imaging of the kinetochore components HCP-4 and ROD-1 showed that no kinetochore is formed during the first embryonic cell division in F2 CENP-A Δ-tail/Δ-tail embryos (Fig 4B, S5-S8 Videos). This kinetochore-null (*knl*) phenotype is also apparent when looking at the speed at which the spindle poles move away from each other following nuclear envelope breakdown (Figs 4C, D; S3 and S4 Videos). Since no physical connection can be established between the chromosomes and microtubules, the lack of tension causes the spindle poles to move away from each other prematurely. We attribute these F2 embryonic phenotypes to the failure in establishing centromere identity in the germ line of F1 homozygotes (CENP-A Δ-tail/Δ-tail).

CENP-A has previously been shown to be dispensable for meiosis. We therefore considered it unlikely that the observed *knl* phenotype in the first mitotic division of F2 CENP-A Δ-tail/Δ-tail embryos is the result of meiotic chromosome segregation defects. Indeed, the meiotic divisions, which happen in the fertilized embryo prior to the onset of the mitotic divisions, appear unaffected (S4 Fig). Therefore, the first instance of observable chromosome segregation defects upon deletion of the CENP-A N-terminal tail appears during the first F2 embryonic division. However, immunostaining of CENP-A and KNL-2 revealed that these proteins have altered localization patterns already in the F1 CENP-A Δ-tail/Δ-tail animals, before the chromosome segregation defects become apparent (S5 Fig). In the mitotic zone of the germ line of F1 CENP-A Δ-tail/Δ-tail animals, both CENP-A and KNL-2 show a punctate pattern, even though mitosis appears to be normal in these cells (S5 Fig). The punctate KNL-2 distribution is also visible in the proximal germ line (S5 Fig). Moreover, the removal of both CENP-A and KNL-2 at the mitosis-to-meiosis transition is delayed, and both proteins can be detected in part of the pachytene region of the germ line.

### *De novo* loading of CENP-A in the *C. elegans* proximal germ line requires KNL-2

CENP-A and KNL-2 have previously been shown to be codependent for chromatin association in embryonic cells [30]. To test the interdependence of CENP-A and KNL-2 in the germ line, we acutely removed these proteins from chromatin in wild type worms by using the AID system with a germline-expressed TIR1 that is present throughout the germ line and in early embryos. Depletion of KNL-2 led to the expected co-depletion of full-length CENP-A in the proximal germ line (Fig 5A). However, depleting full-length CENP-A in the proximal germ line did not lead to loss of KNL-2, as was reported in embryos. Instead, KNL-2 remained largely associated with chromatin (Fig 5A). The persistence of KNL-2 in proximal germ cells upon depletion of full-length CENP-A suggests that KNL-2 acts upstream of CENP-A in the *de novo* establishment of centromeres, which may explain the essential role of the CENP-A N-terminal tail in this process. Consistently, CENP-A Δ-tail remains visible on chromatin upon depletion of KNL-2 in F1 CENP-A Δ-tail homozygotes, suggesting that its chromatin association is no longer dependent on the presence of KNL-2 (Fig 5B).

**Fig 5.**
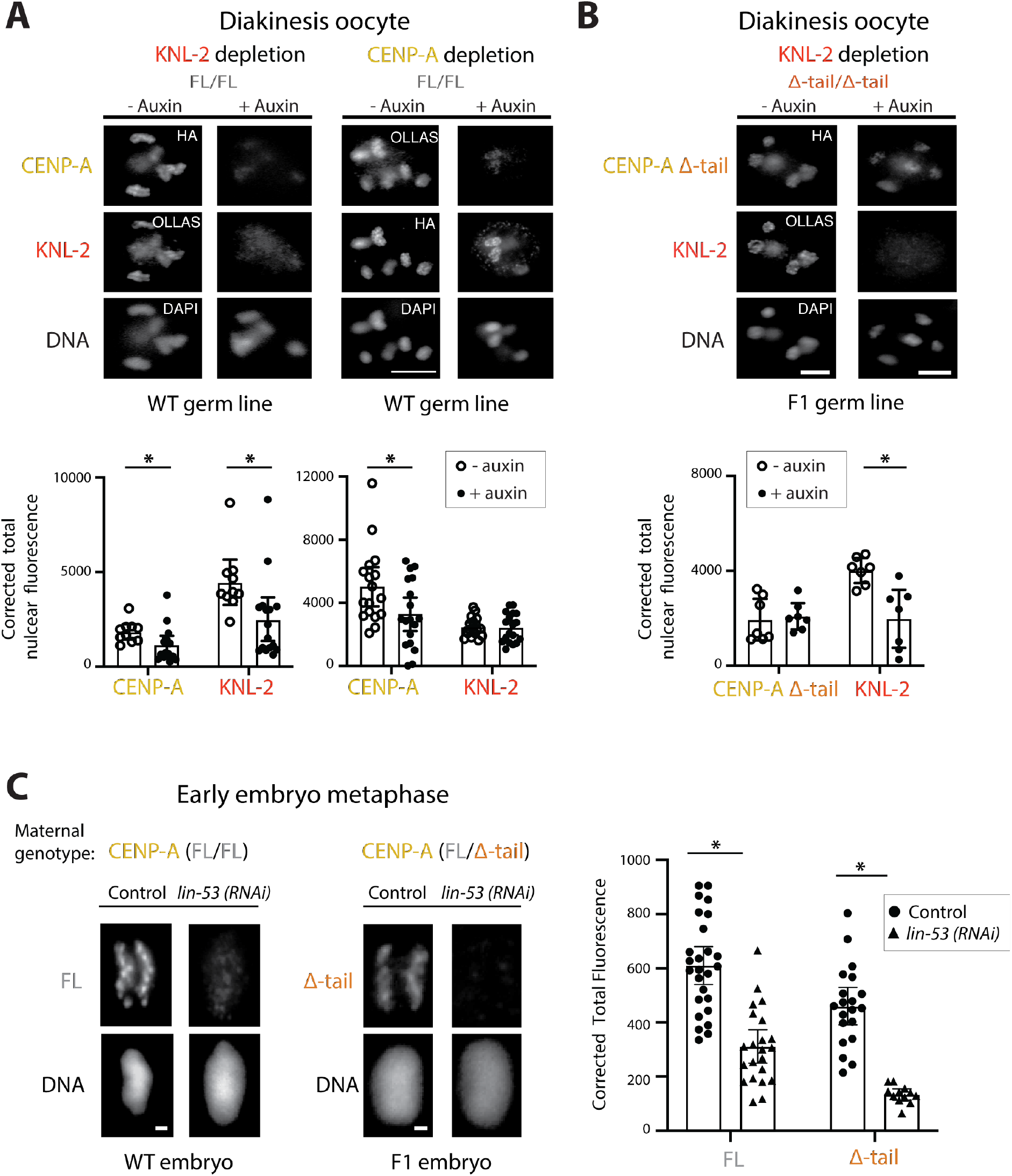
Chromatin association of CENP-A Δ-tail depends on LIN-53 and not KNL-2. (A) Germline-specific depletion of CENP-A and KNL-2 using the auxin-induced degradation system. IF images of CENP-A and KNL-2 in diakinesis oocytes (−2 position) of adult hermaphrodite germ lines, with and without auxin-induced degradation of either KNL-2 (left) or CENP-A (right). Bar plots show quantifications of total nuclear fluorescence in diakinesis oocytes (−1 to −3 positions). (B) Chromatin association of CENP-A Δ-tail in presence or absence of KNL-2. IF images and fluorescence quantifications of CENP-A Δ-tail and KNL-2 in diakinesis nuclei of CENP-A Δ-tail F1 homozygous adult hermaphrodite germ lines. KNL-2 was depleted using germline-specific AID. (C) Chromatin association of CENP-A Δ-tail in presence or absence of LIN-53. IF images showing CENP-A FL and CENP-A Δ-tail at metaphase in early embryos. *Lin-53* was depleted by RNA interference. Bar plots show quantifications of CENP-A levels at metaphase. Germline-specific depletion of KNL-2 and CENP-A in (A) and (B) was achieved using a TIR1 with germline- and early embryo-restricted expression. In (A-C), error bars show the 95% confidence interval of the mean, and the asterisks denote a statistically significant difference, determined by using the’s t-test. Scale bars represent 5 μm (A, B) or 1 μm (C).

### CENP-A is maintained on chromatin in absence of the interaction with KNL-2

The chromatin association of CENP-A Δ-tail in the F1 homozygotes seems puzzling, given the observations that KNL-2 is required for the chromatin association of full-length CENP-A and that the CENP-A N-terminal tail is required for the interaction with KNL-2. It implies that the maintenance of CENP-A Δ-tail during development of the F1 CENP-A Δ-tail/Δ-tail animals is mediated by factors other than KNL-2. The RbAp46/48 homologue LIN-53 has recently been shown to be important for CENP-A chromatin-association in *C. elegans* [31]. We found that depleting LIN-53 by RNAi resulted in reduced levels of both full-length CENP-A and the CENP-A Δ-tail, suggesting that LIN-53 could be involved in ensuring centromere maintenance in the homozygous CENP-A Δ-tail F1 animals (Fig 5C).

In summary, we propose a functional distinction between centromere establishment and centromere maintenance in *C. elegans*. Though both require CENP-A, the CENP-A N-terminal tail is specifically required to re-establish centromeres in the proximal adult hermaphrodite germ line in every generation, and mediates an interaction with KNL-2. Once established, functional centromeres are inherited and maintained in absence of the N-terminal tail, but only until the next generation, where resetting of centromere identity fails (Fig 6A).

**Fig 6.**
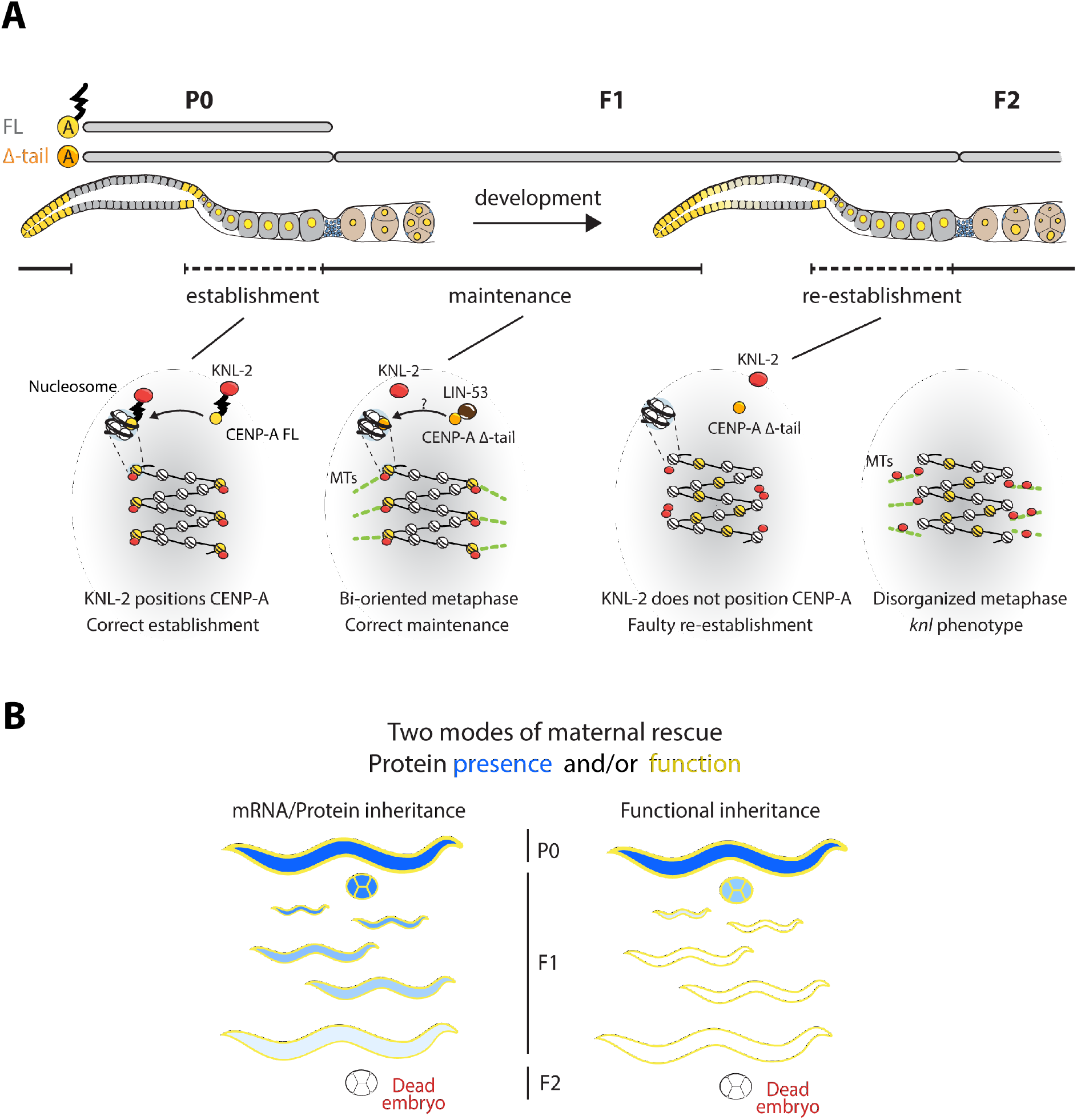
The CENP-A N-terminal tail is required for the establishment of centromere identity that is reset in every generation. (A) Model of centromere establishment and maintenance across generations in *C. elegans*. Centromeres are established in the proximal germ line and maintained until their removal in the distal germ line of the next generation. The CENP-A N-terminal tail interacts with KNL-2 and is required for establishing centromeres. In the heterozygous P0 worm, the CENP-A full-length present in the germ line is sufficient for this process. Upon loss of the CENP-A N-terminal tail, centromeres can be maintained for one generation, a process that likely involves the histone chaperone LIN-53. However, in absence of the CENP-A N-terminal tail, centromere re-establishment fails in the adult F1 generation, leading to a mislocalization of CENP-A, KNL-2 and a kinetochore-null phenotype in the F2 embryos. (B) Models for explaining a maternal-effect lethal phenotype, where the function of an essential gene is maintained for one generation despite loss or truncation of the gene. Left, sufficient protein or mRNA is deposited in the embryo from the maternal germ line for it to persist and function during development. Right, the protein sets a (chromatin) state in the parental germ line that serves as a memory that persists during development and allows its function to be retained. Protein presence is labeled in blue, and function in yellow.

## Discussion

In this study, we take advantage of the CENP-A dynamics in the *C. elegans* adult hermaphrodite germ line to investigate the regulation of centromere formation and inheritance. In most eukaryotes, CENP-A is maintained by an epigenetic mechanism from one cell cycle to the next. DNA replication dilutes CENP-A nucleosomes at established functional centromeres, and the remaining CENP-A nucleosomes act as epigenetic marks for the re-loading of new CENP-A at the same genomic regions at cell cycle stages that vary between organisms [10,38–40]. CENP-A is also maintained from one generation to the next in many model organisms. CENP-A molecules in mouse oocytes show remarkable stability during mouse meiosis, a feature that is dependent on the MIS18 complex [25]. In contrast, CENP-A in starfish oocytes is actively replenished at centromeres [24]. This mechanism is again dependent on the MIS18 complex, as well as the chaperone HJURP, and active transcription to remove the old CENP-A-containing nucleosome.

In *C. elegans*, the cycle of inheritance is interrupted at the onset of meiosis in hermaphrodites, and centromeres have to be re-established during diplotene/diakinesis of meiosis I in the proximal germ line (Fig 1). How centromeres are established *de novo* is not well understood, and most studies so far have relied on the observation of spontaneous or artificially induced neocentromeres or artificial chromosomes. The factors required for this process are similar to those required for centromere maintenance. HJURP, CENP-C and CENP-I have all been implicated in centromere establishment by studying the formation of neocentromeres in human and chicken cell lines [41–43].

The MIS18 complex or M18BP1/KNL-2 has not been reported to be sufficient for neocentromere establishment, as this complex is believed to only recognize existing centromeres through interactions with CENP-C or CENP-A [44,45]. However, non-mammalian KNL-2s often show cell cycle dynamics that are different from their human homologue, and localize to centromeres throughout the cell cycle, suggesting that they may have taken on additional roles [30,44–47]. Consistently, we find in this study that *C. elegans* KNL-2 is essential both for centromere establishment and maintenance. However, its role may be different during the two processes. CENP-A and KNL-2 are codependent for chromatin association during maintenance, as previously shown [30], while we find that KNL-2 may act upstream during establishment.

Our results show that the interaction between KNL-2 and CENP-A depends on the central region of KNL-2 and the N-terminal tail of CENP-A in *C. elegans*. In chicken cells, the interaction between KNL-2 and CENP-A was shown to be dependent on a KNL-2 CENP-C motif [45]. Such a CENP-C motif has been identified in a wide variety of taxa, including fish, frogs and plants [46,48,49]. In *A. thaliana*, disrupting the CENP-C motif leads to a failure of KNL-2 to bind to centromeric chromatin [49]. Additionally, a detailed study of *Xenopus laevis* KNL-2 shows that the CENP-C motif regulates interaction with CENP-A nucleosomes [46]. Considering the conserved nature of the CENP-C motif, it is perhaps surprising that it cannot be found in *C. elegans* KNL-2 [50]. Instead, the minimal region for interaction with full-length CENP-A in yeast two-hybrid had already been shown to be a larger central region of KNL-2 (aa 269-470) [33]. We could confirm this result and demonstrate that *in vitro* the interaction is mediated by the N-terminal tail of CENP-A. Moreover, we show that the CENP-A N-terminal tail is both necessary and sufficient for the interaction with KNL-2 *in vivo*.

The N-terminal tail of CENP-A is considered to be flexible and unstructured, but plays an important role in centromere biology [51–55]. It contributes to the interactions with the kinetochore proteins CENP-B, CENP-C and CENP-T [51,52,56,57], although there may be species-specific differences in how these proteins interact. Different post-translational modifications (PTMs) on the CENP-A N-terminal tail have also been described to affect centromere establishment and maintenance [53,54,58–63].

Here, we describe a role for the CENP-A N-terminal tail in establishing centromere identity in the hermaphrodite germ line of *C. elegans*. In *A. thaliana*, the CENP-A N-terminal tail is required for meiosis, and CENP-A N-terminal tail replacements caused defects in meiotic CENP-A loading, defects in meiotic segregation, and led to sterility [64,65]. In fission yeast, removing the CENP-A N-terminal tail does not affect the loading of CENP-A at centromeres [57]. However, the N-terminal tail does confer centromere stability in mitotically dividing cells, most likely through an epigenetic process [57]. We propose that in *C. elegans* the CENP-A N-terminal tail has a role during meiosis, though not in supporting meiotic divisions, since CENP-A is dispensable in this context. However, it is required to interact with KNL-2 and to define centromere identity, perhaps similar to what was observed in fission yeast [57], which allows centromere inheritance to the next generation.

We report that CENP-A lacking the N-terminal tail is no longer dependent on KNL-2 for chromatin association. Strikingly, this truncated version of CENP-A is still functional in mitotic chromosome segregation, and we therefore hypothesize that KNL-2 has essential functions in mitosis other than CENP-A loading. LIN-53, a homologue of the histone chaperone RbAp46/48, has been implicated in maintaining CENP-A chromatin association in *C. elegans* [31]. We find that this is the case also for CENP-A lacking the N-terminal tail, implicating LIN-53 in centromere maintenance, although we have not ruled out an additional role in centromere establishment.

While the first generation homozygous for the CENP-A N-terminal tail deletion (F1) is viable, and CENP-A Δ-tail is sufficient to properly assemble the kinetochore, the F2 embryos show the characteristic features of a kinetochore-null phenotype [32,66,67]. F1 and F2 homozygous embryos are genetically identical, and we therefore propose that the kinetochore defects in F2 embryos do not stem from defects in recruiting specific components, but rather from the fact that centromere identity has not been established in the maternal germ line of the previous generation. A question that remains is when centromere identity is determined. We speculate this happens when CENP-A is re-loaded onto chromatin at the diplotene stage of the proximal germ line (Fig 1). However, it is also possible that it happens after fertilization, since CENP-A localization appears to be dynamic during the meiotic divisions [28].

The CENP-A Δ-tail strain shows a maternal effect lethal phenotype. This phenotype is often explained by a maternal supply of messenger RNA (mRNA) or proteins into the embryo that is sufficient to fulfill the essential functions during development (Fig 6B). In *D. melanogaster* many maternal effect genes affect the development of the early embryo through spatially defined depositioning of maternal RNAs or protein [68]. In *C. elegans*, many genes have been reported to show a maternal effect when mutated [69–73]. A recent study in this species also showed that maternal proteins and RNA can be inherited by a null mutant until the first larval stage [74]. We ruled out that the viability of the CENP-A Δ-tail F1 homozygotes solely depends on the presence of the maternal full-length protein. Instead, it is dependent on the presence of the full-length protein in the germ line of the preceding generation (P0). We therefore hypothesize that information about centromere identity rather than full-length CENP-A protein is transferred from the maternal germ line into the zygote (Fig 6B). This form of inheritance has been reported previously in *C. elegans*, often for proteins that stabilize chromatin marks [75]. One example is the polycomb repressive complex 2 (PRC2), which maintains H3K27me3 during embryogenesis [76,77]. Recently, the H3.3 chaperone HIRA has also been implicated in setting a heritable chromatin state in the maternal germ line that can be maintained into adulthood [78]. Further work will be required to uncover how the interaction between KNL-2 and the CENP-A N-terminal tail can establish a chromatin state that defines the centromere for an entire generation.

## Materials and Methods

### Nematode maintenance and strain generation

*C. elegans* strains were maintained using standard conditions at 20°C unless otherwise noted. N2 was used as a wild type strain and all of the genetic modifications were performed in this background unless otherwise indicated. The hT2 [bli-4(e937) let-?(q782) qIs48] (I;III) balancer allele was used to maintain the *hcp-3* (CENP-A) deletion and truncation alleles. Heterozygote adults of these strains give rise to 4/16 heterozygous *hcp-3*, 1/16 homozygous *hcp-3*, and 11/16 inviable hT2 aneuploid or hT2 homozygous offspring. S1 Table provides details for the strains used in this study. All insertions and deletions were generated using CRISPR/Cas9 technology as described in [79]. RNAi experiments were performed using clones from the Ahringer library [80].

### Immunofluorescence microscopy

For immunofluorescence experiments in the germ line and embryos, young gravid adult worms were dissected in anaesthetizing buffer (50 mM sucrose, 75 mM HEPES pH 6.5, 60 mM NaCl, 5 mM KCl, 2 mM MgCl_2_, 10 mM EGTA pH 7.5, 0.1% NaN_3_). The dissected germ lines and embryos were freeze-cracked and fixed in methanol for 5 min at −20 °C. Slides were washed with PBS and incubated with anti-HA antibody (mouse, clone 42F13, 1:60), anti-OLLAS antibody (rat, Novus Biologicals NBP1-06713, 1:200), or anti-α-Tubulin (mouse, Sigma-Aldrich T9026, 1:1000) overnight at 4 °C. Slides were washed in PBS + 0.05% Tween and incubated with Cy3- or Alexa 488-conjugated secondary antibodies (Jackson Immunoresearch, 1:700; Invitrogen, 1:1000) for 1.5 h at room temperature. Samples were washed with PBS + 0.05% Tween, stained with DAPI, and mounted with VECTASHIELD Antifade Mounting Medium. Images were obtained using a Leica DM5000 B microscope.

### Live imaging

Young gravid adults were dissected on a 2% agarose pad and kept in egg buffer (118mM NaCl, 48 mM KCL, 2mM CaCl2, 2mM MgCl2 and 25mM Hepes, pH 7,5). All images were acquired using the Intelligent Imaging Innovations Marianas SDC spinning disk system, mounted on an inverted microscope (Leica DMI 6000B). Embryos were kept at 20°C, and imaged with a 63x glycerin objective (NA=1.4), and an emCCD Evolve 512 camera (Photometrics). Images were captured in stacks of 10 sections along the z axis at 1-μm intervals every 10 s with 100-200 ms exposures (depending on the strain) at the 488-nm (100%) and 594-nm (100%) channels. Nuclear envelope breakdown (NEBD) was visually determined and used as point zero in time series analysis.

### Image and statistical analysis

All images were processed using Fiji software [81]. Z-stack and single plane images were adjusted for contrast and brightness. To assess the intensity of immunofluorescence stainings or western blot bands, corrected total fluorescence (CTF) was calculated using the following formula: CTF = Integrated Density – (Area x Mean fluorescence of background). The significance of observed differences was tested for by using the Student’s t-test.

### Co-immunoprecipitation and western blotting

Synchronized young gravid adult worms were washed three times with M9, and embryos were obtained by hypochlorite treatment. Embryos were resuspended in lysis buffer (50 mM Tris-HCl pH = 7.4, 500 mM NaCl, 0.25% deoxycholate, 10% glycerol, 1% NP-40) and frozen in liquid nitrogen. Samples were sonicated with a Bioruptor Pico (Diagenode - 15 cycles, 30 s sonication, 30 s rest, snap freezing between each cycle) and spun down (30 min, max speed) to pellet debris. The supernatant was collected, and pre-washed Pierce™ Anti-HA Magnetic Beads (Thermo Fisher Scientific) were added. Following overnight incubation on a rotator at 4°C, the beads were collected using a magnetic stand and washed according to manufacturer’s instructions. HA-tagged proteins were eluted by boiling the beads for 5-10 minutes in Pierce™ Lane Marker Non-Reducing Sample Buffer (Thermo Fisher Scientific). Protein concentrations in input samples were measured with the Bio-Rad Protein Assay (Bio-Rad, 5000006) and equal amounts were loaded on SDS-page gels. Western blotting was performed according to standard procedures using the LI-COR Odyssey system and 4-20% gradient gels. Primary antibodies (anti-HA, Sigma-Aldrich mAb 3F10, 1:1000; anti-OLLAS, Novus Biologicals NBP1-06713B, 1:1000) were incubated overnight at 4 °C, and IRDye^®^ secondary antibodies appropriate for each primary antibody were incubated for 1 h at room temperature.

### Yeast two-hybrid experiments

The yeast two-hybrid assay was performed using the GAL4-based system as previously described by [82]. In short, the yeast strain MAV203 was transformed according to the manufacturer’s instructions with pDEST22 (Invitrogen) plasmids expressing transcriptional activation domain fusions, or pDEST32 (Invitrogen) plasmids expressing DNA binding domain fusions. The KNL-2 cDNA fragments were cloned into pDEST22 and CENP-A cDNA fragments into pDEST32. The clones were spotted on non-selective medium (leucine and tryptophan double dropout) and on selective medium (leucine, tryptophan and histidine triple dropout with 5 mM 3AT (3-amino-1,2,4-triazole)).

### Auxin-induced degradation (AID)

Strains for auxin-induced degradation were constructed by introducing a degron tag at the gene locus of interest. These strains were crossed to the germline- or soma-specific TIR1 drivers described in [37], and auxin treatment was performed by transferring worms to bacteria-seeded plates containing 4 mM natural auxin (indole-3-acetic acid; Alfa Aesar). Worms were kept on these plates for 4 h (germline depletion) or 3 d (somatic depletion) at 20°C.

### *In vitro* purification of recombinant proteins and interaction assay

The nucleotide sequence corresponding to KNL-2 aa 269-470 (central region) and CENP-A aa 1-179 (N-terminal tail) were cloned into a vector derived from pET for expression in *E. coli* as fusion proteins containing for KNL-2 a N-terminal His_9_-MBP tag with a TEV cleavage site and for CENP-A an N-terminal His_9_-MBP tag with a TEV cleavage site and a C-terminal GST tag. The recombinant proteins were overexpressed in *E. coli* BL21-Star cells, grown in TB media at 37°C for 6 h followed by overnight induction at 18°C with 0.1 mM isopropyl-β-D-thiogalactopyranoside (IPTG). His_9_-MBP-TEV-KNL2^269-470^ induced cells were harvested by centrifugation and resuspended in TBS buffer for MBP purification (50mM Tris-Cl pH 8.0, 750mM NaCl, 1mM DTT, 0,15% CHAPS, 1 μg/ml DNase, 1 μg/ml Lysozyme), supplemented with protease inhibitors (1mM PMSF, 1 μg/ml leupeptin, and 2 μg/ml pepstatin). His_9_-MBP-TEV-CENP-A^1-179^-GST induced cells were harvested by centrifugation and resuspended in lysis/washing buffer for NiNTA purification (50 mM phosphate buffer pH 8.0, 300mM NaCl, 25mM imidazole, 0.15% CHAPS, 5 mM β-mercaptoethanol, 1 μg/ml DNase, 1 μg/ml Lysozyme), supplemented with protease inhibitors. The control GST peptide was purified from the pET42 empty vector using the same conditions as described for CENP-A^1-179^. Cells were lysed using an emulsiflex system (AVESTIN). The soluble fraction was subjected to an affinity purification using a MBP FF crude column (GE Healthcare) for the KNL-2 fusion construct, or a chelating HiTrap FF crude column (GE Healthcare) charged with Ni^2+^ ions for the CENP-A fusion constructs and the GST control. The proteins were eluted and desalted on a desalting column (GE Healthcare). For the *in vitro* interaction assay, the KNL-2 and CENP-A recombinant proteins were mixed in equimolar amounts. GST was used as a negative control. After 1 hour incubation at RT, Tobacco Etch Virus protease (TEV) was added and digestion was performed 30 min at RT. Glutathione High Capacity Magnetic Agarose Beads (Sigma G0924) were added and the mixture was rotated 1 hour at RT. The beads were then washed and eluted, after which all fractions were loaded onto a 4%-20% gradient SDS-PAGE gel and stained with Coomassie Brilliant Blue after electrophoresis.

## Acknowledgements

We thank members of the Steiner lab for help and discussion, Dario Menéndez for help with strain generation, Marina Berti for reagent preparation, Nicolas Roggli for help with the figure preparation, Patrick Meraldi and Nikolai Klena for comments on the manuscript, and Stella Toonen for proofreading. Some strains were provided by the CGC, which is funded by NIH Office of Research Infrastructure Programs (P40 OD010440). We are grateful to the Bioimaging Center of the Faculty of Sciences at the University of Geneva.

## Author contributions

FAS and RFP conceived the study. RFP carried out all genetics and immunofluorescence experiments, JMW carried out live cell imaging experiments, CG carried out the *in vitro* interaction experiments, KD did the majority of the CRISPR injections for strain generation, RFP, FS and MG carried out the yeast two-hybrid experiments in the laboratory of MG. RFP and FAS prepared the manuscript with input from the other authors. FAS acquired funding.

---

**S1 Fig.**
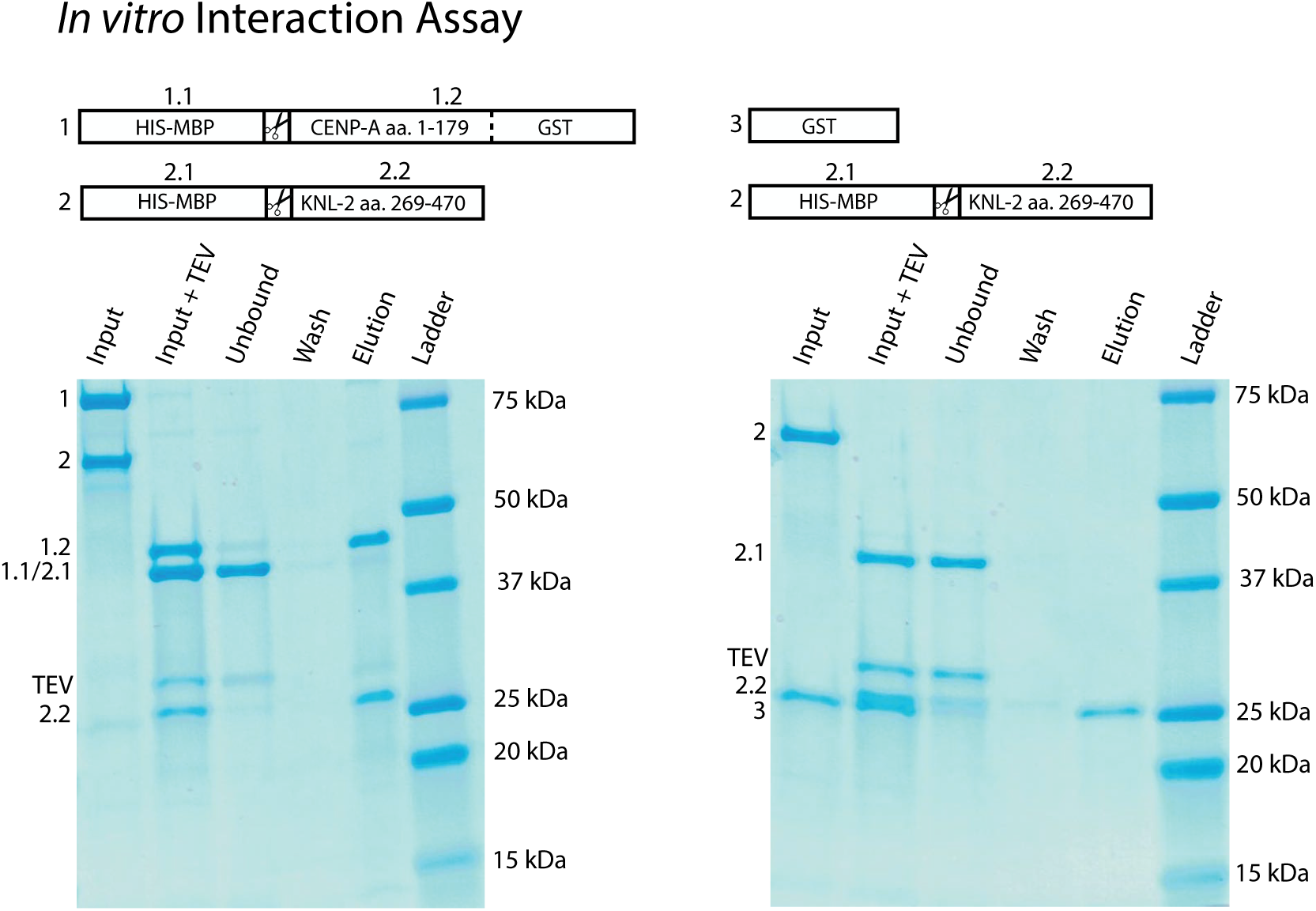
The CENP-A N-terminal tail interacts with the KNL-2 central domain *in vitro*. *In vitro* interaction of CENP-A N-terminal tail and KNL-2 central domain. CENP-A N-terminal tail (aa 1-179) fused to GST, GST alone, and the KNL-2 central region (aa 269-470) were purified from bacteria. To aid solubility, a HIS-MBP tag was fused to the KNL-2 and CENP-A peptides. The MBP tag was removed by TEV cleavage before the pull-down using GST beads. The GST::CENP-A N-terminal tail (left) co-precipitates the KNL-2 central fragment, while GST alone (right) does not.

**S2 Fig.**
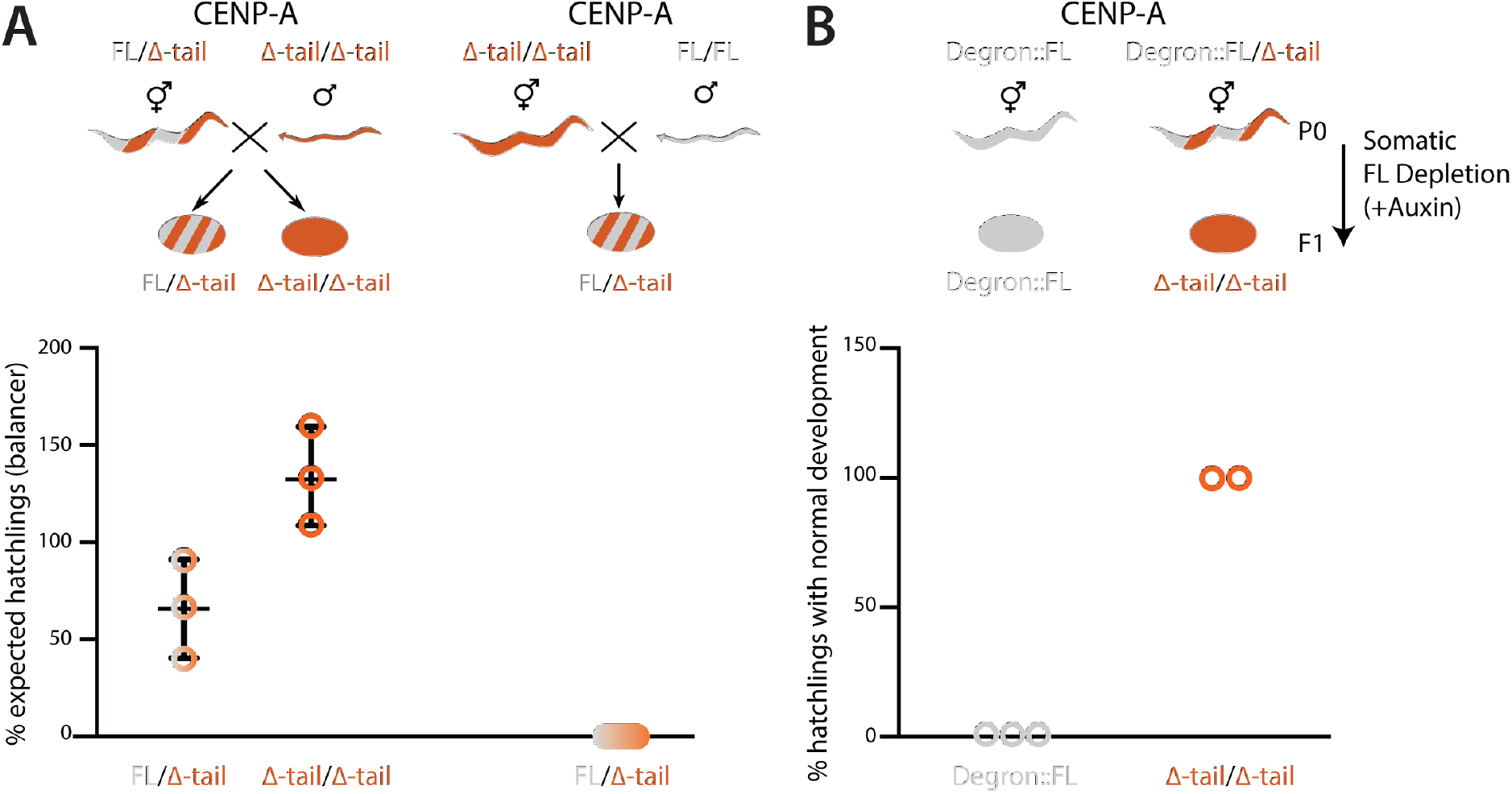
Full-length CENP-A is deposited maternally, but the deposited pool does not persist through development. Percentage of viable cross-progeny after crossing worms carrying CENP-A FL (gray) and Δ-tail (orange) alleles. Left, crossing homozygous CENP-A Δ-tail males with heterozygous FL/Δ-tail hermaphrodites gives rise to fully viable progeny, showing that the homozygous Δ-tail males are fertile. Right, crossing wild type males to homozygous CENP-A Δ-tail hermaphrodites does not rescue the embryonic lethality. Error bar shows the standard deviation of the mean. (B) Quantification of hatchlings developing normally after somatic AID-mediated depletion of CENP-A FL during development in worms homozygous for CENP-A FL or Δ-tail. Worms only carrying the degron-tagged CENP-A FL protein are inviable, whereas worms homozygous for the CENP-A Δ-tail are unaffected by this treatment.

**S3 Fig.**
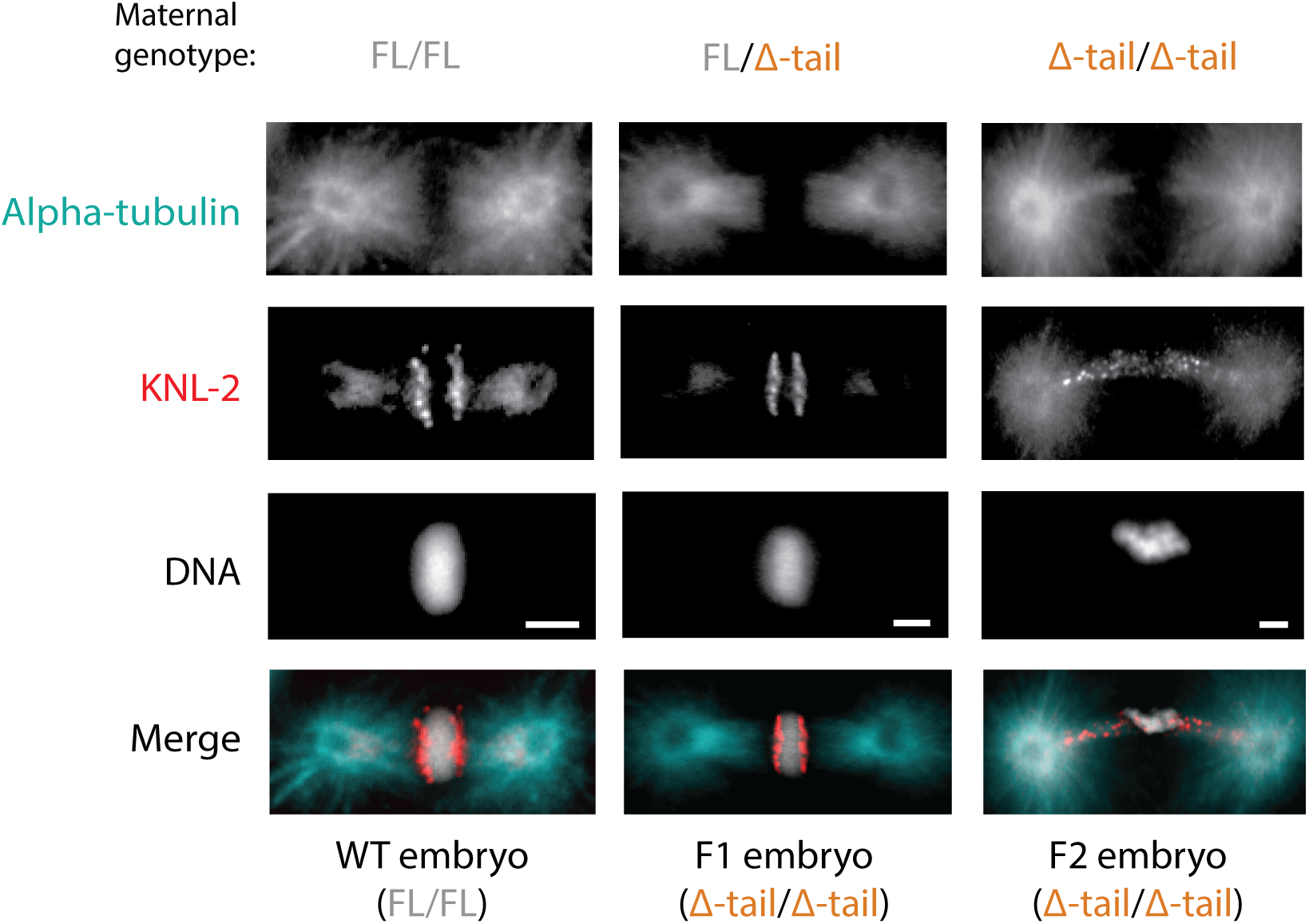
KNL-2 accumulates on microtubules during metaphase in F2 homozygous CENP-A Δ-tail embryos. IF images depicting KNL-2 and α-tubulin at metaphase of early embryonic cell divisions in embryos derived from CENP-A FL/FL, FL/Δ-tail or Δ-tail/Δ-tail maternal germ lines. Scale bars represent 2 μm.

**S4 Fig.**
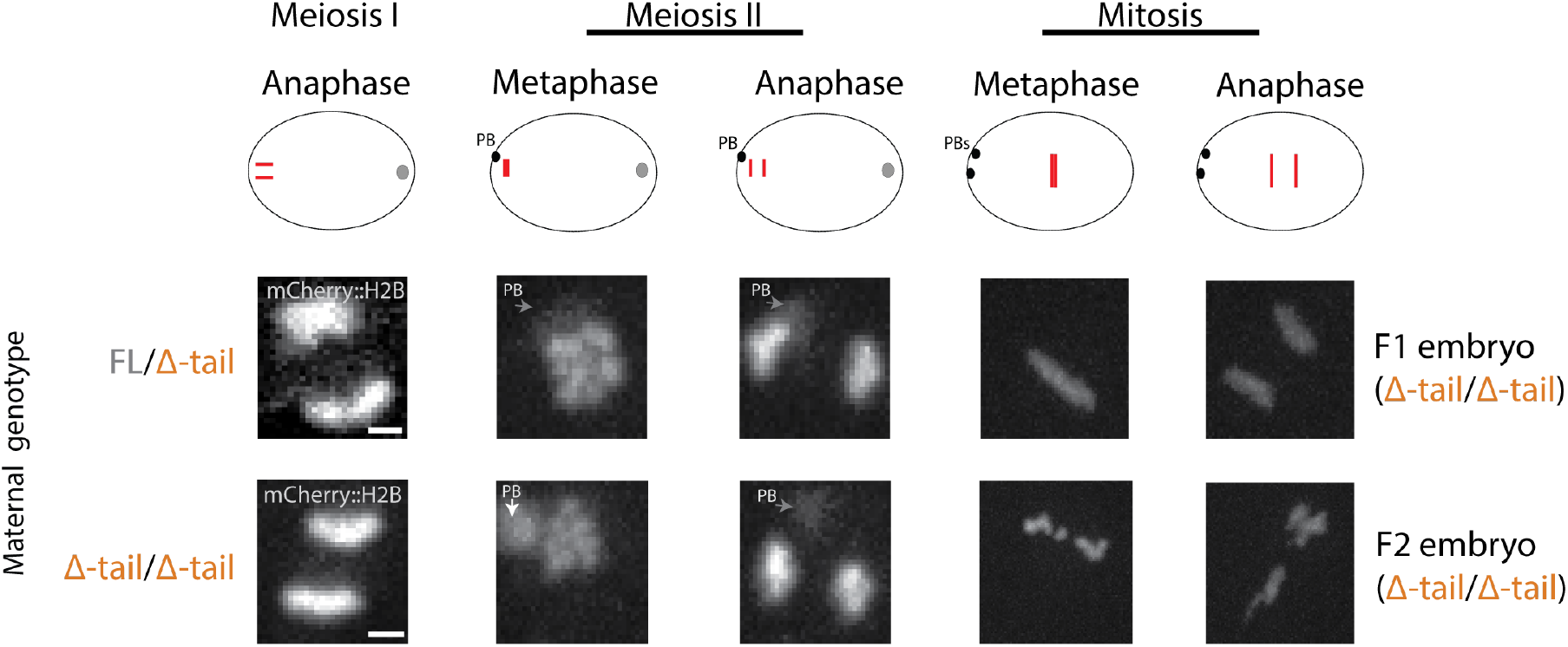
Meiotic segregations are unaffected in CENP-A Δ-tail homozygotes. Still images of live cell recordings of embryos derived from CENP-A FL/Δ-tail or Δ-tail/Δ-tail maternal germ lines. H2B::mCherry was used to visualize chromosome segregation in meiosis I and meiosis II, and mitosis of the first embryonic cell division. Polar bodies are labeled with PB and arrows. Cartoon images of embryos with the chromosomes corresponding to the images in red. Scale bars represent 2 μm.

**S5 Fig.**
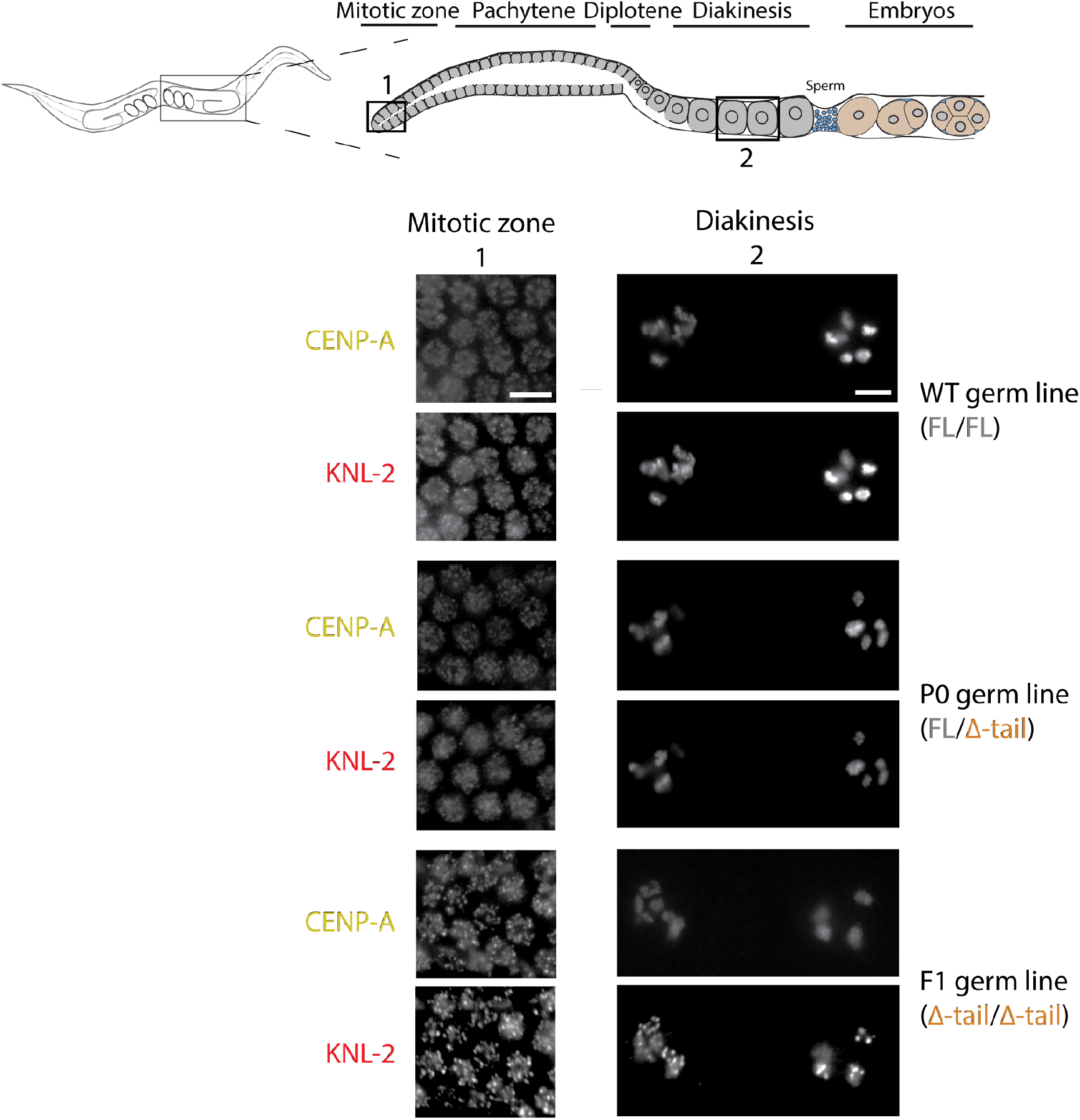
CENP-A and KNL-2 patterns in the germ line of worms with different CENP-A genotypes. IF images showing the patterns of KNL-2 and CENP-A FL and Δ-tail in distal mitotic and diakinesis nuclei, as highlighted in the cartoon image of the germ line, in the adult germ line of CENP-A FL/FL (wild type control), FL/Δ-tail (P0) and Δ-tail/Δ-tail (F1) animals. Scale bars represent 5 μm.

**S1 Video. KNL-2::GFP in first embryonic division for embryos derived from CENP-A FL/Δ-tail maternal germ line (F1 embryo).**

**S2 Video. KNL-2::GFP in first embryonic division for embryos derived from CENP-A Δ-tail/Δ-tail maternal germ line (F2 embryo).**

**S3 Video. mCherry::H2B and mCherry::γ-tubulin in first embryonic division for embryos derived from CENP-A FL/Δ-tail maternal germ line (F1 embryo).**

**S4 Video. mCherry::H2B and mCherry::γ-tubulin in first embryonic division for embryos derived from CENP-A Δ-tail/Δ-tail maternal germ line (F2 embryo).**

**S5 Video. ROD-1::GFP in first embryonic division for embryos derived from CENP-A FL/Δ-tail maternal germ line (F1 embryo).**

**S6 Video. ROD-1::GFP in first embryonic division for embryos derived from CENP-A Δ-tail/Δ-tail maternal germ line (F2 embryo).**

**S7 Video. HCP-4::mCherry in first embryonic division for embryos derived from CENP-A FL/Δ-tail maternal germ line (F1 embryo).**

**S8 Video. HCP-4::mCherry in first embryonic division for embryos derived from CENP-A Δ-tail/Δ-tail maternal germ line (F2 embryo).**

**S1 Table.**
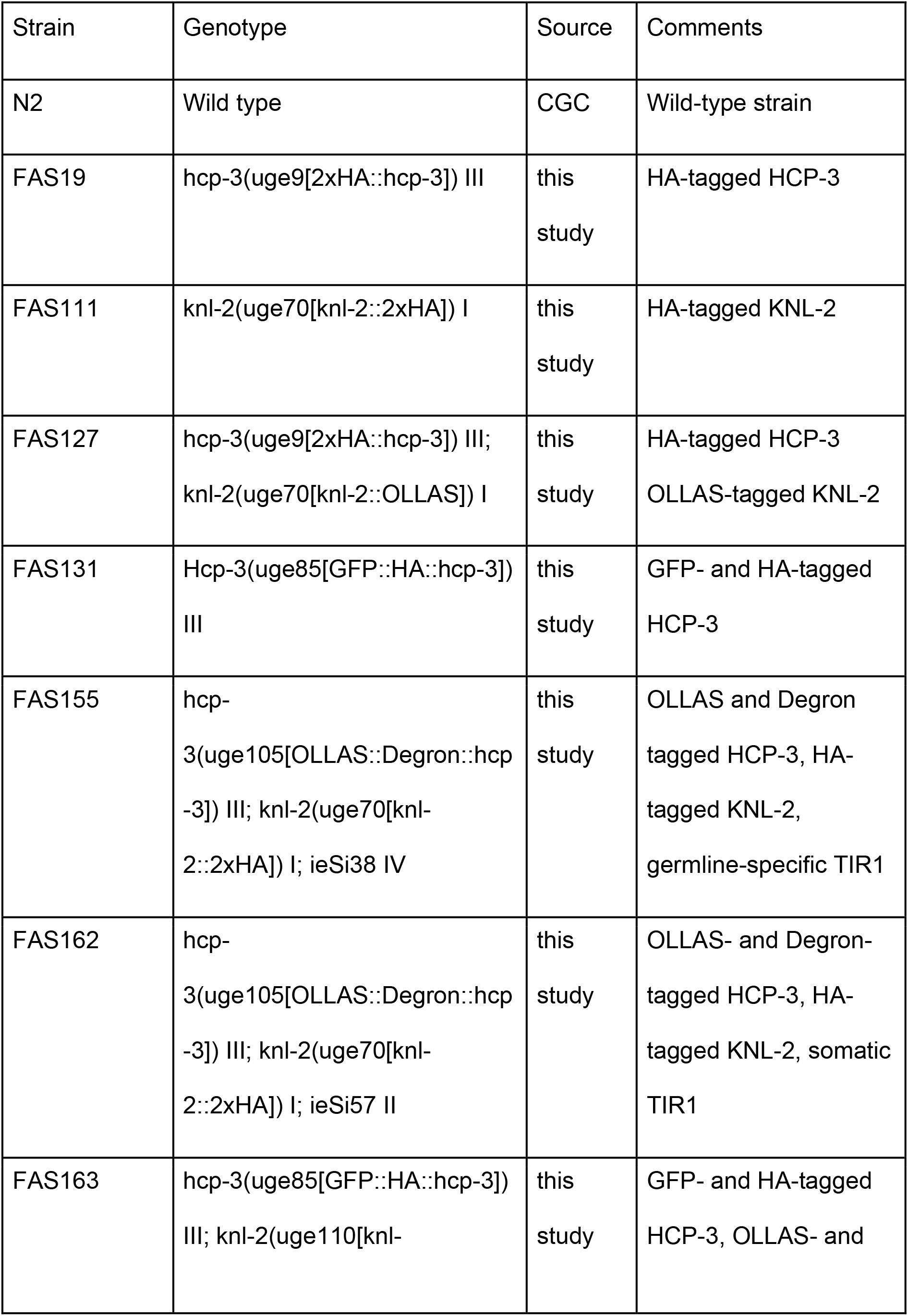
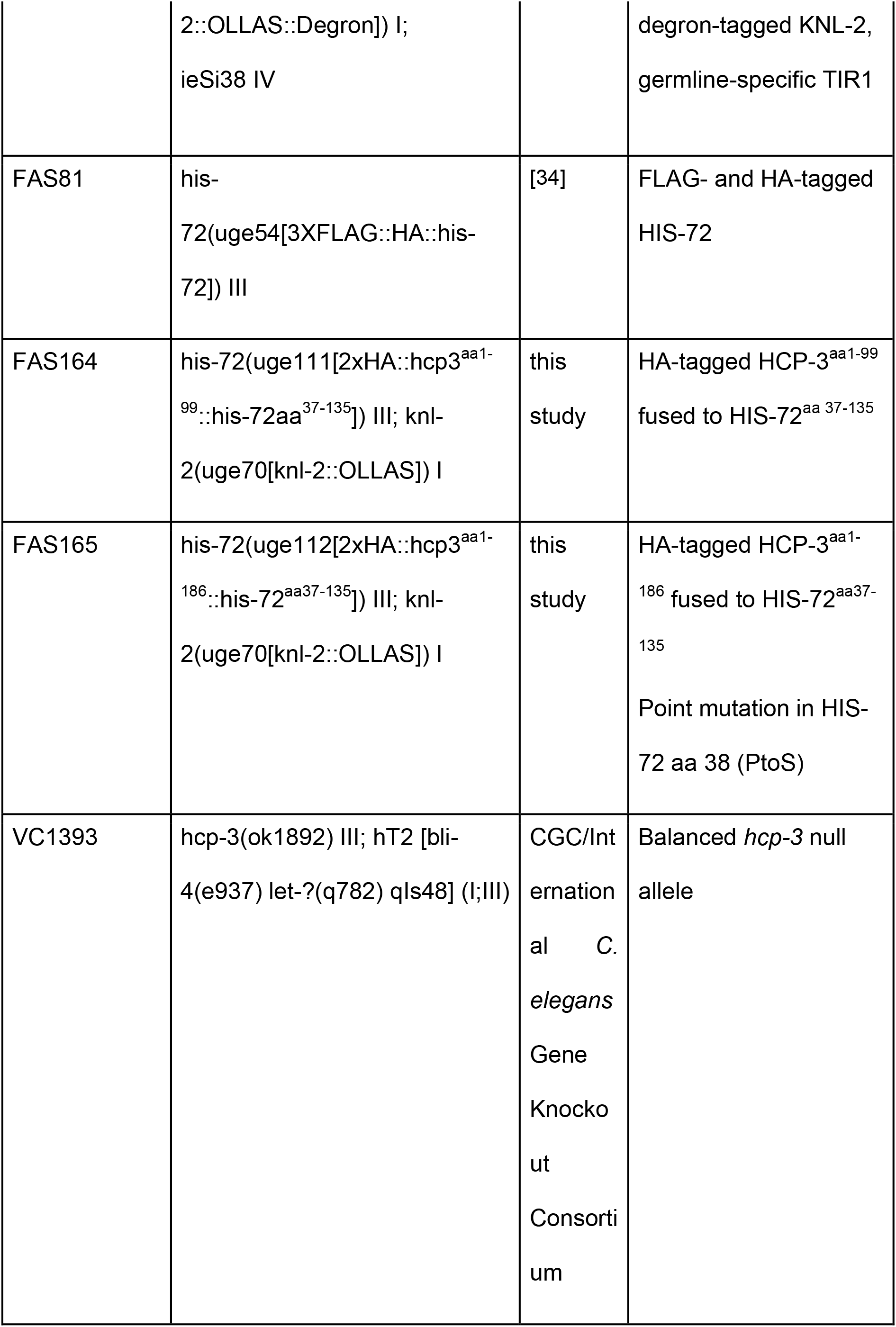
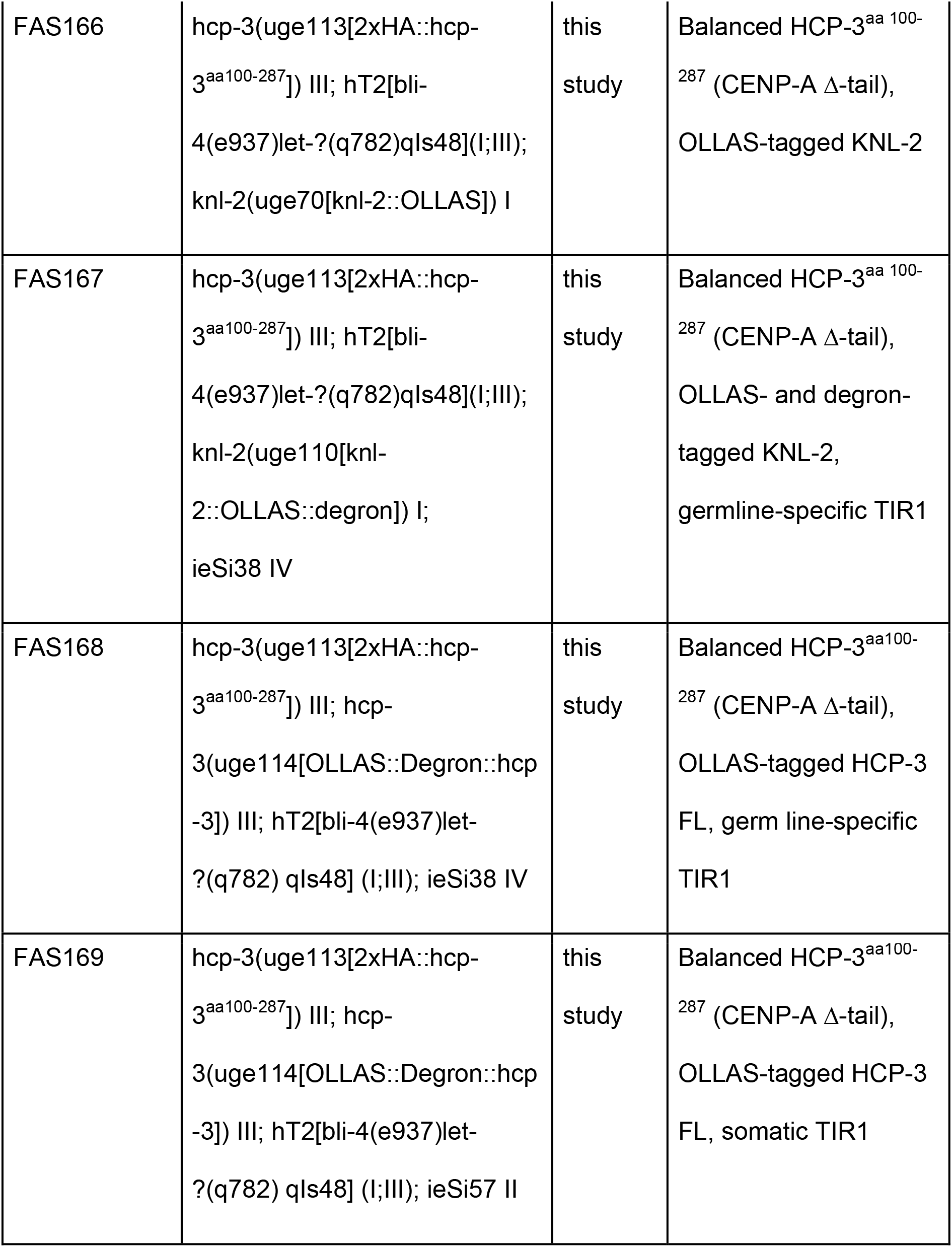
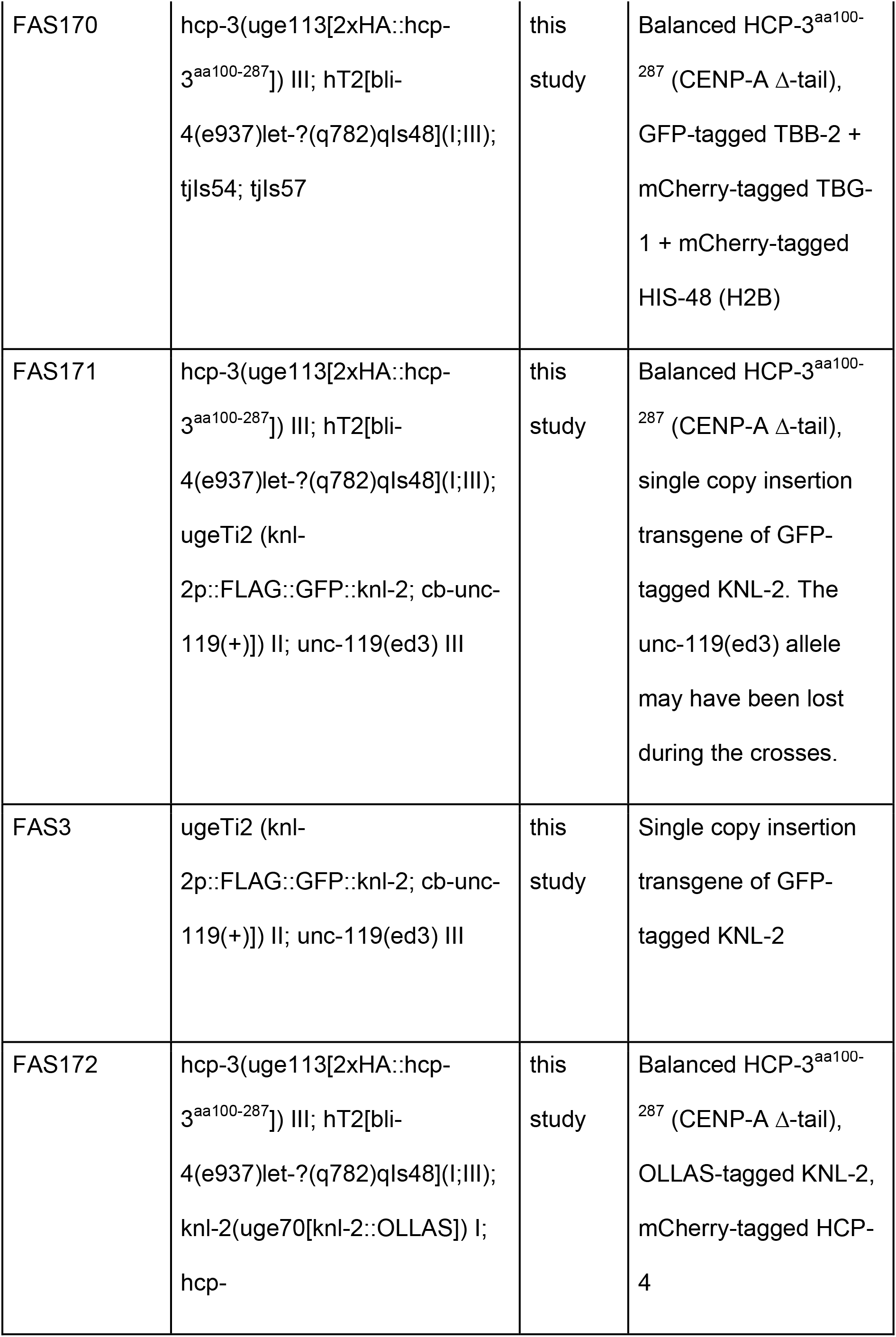
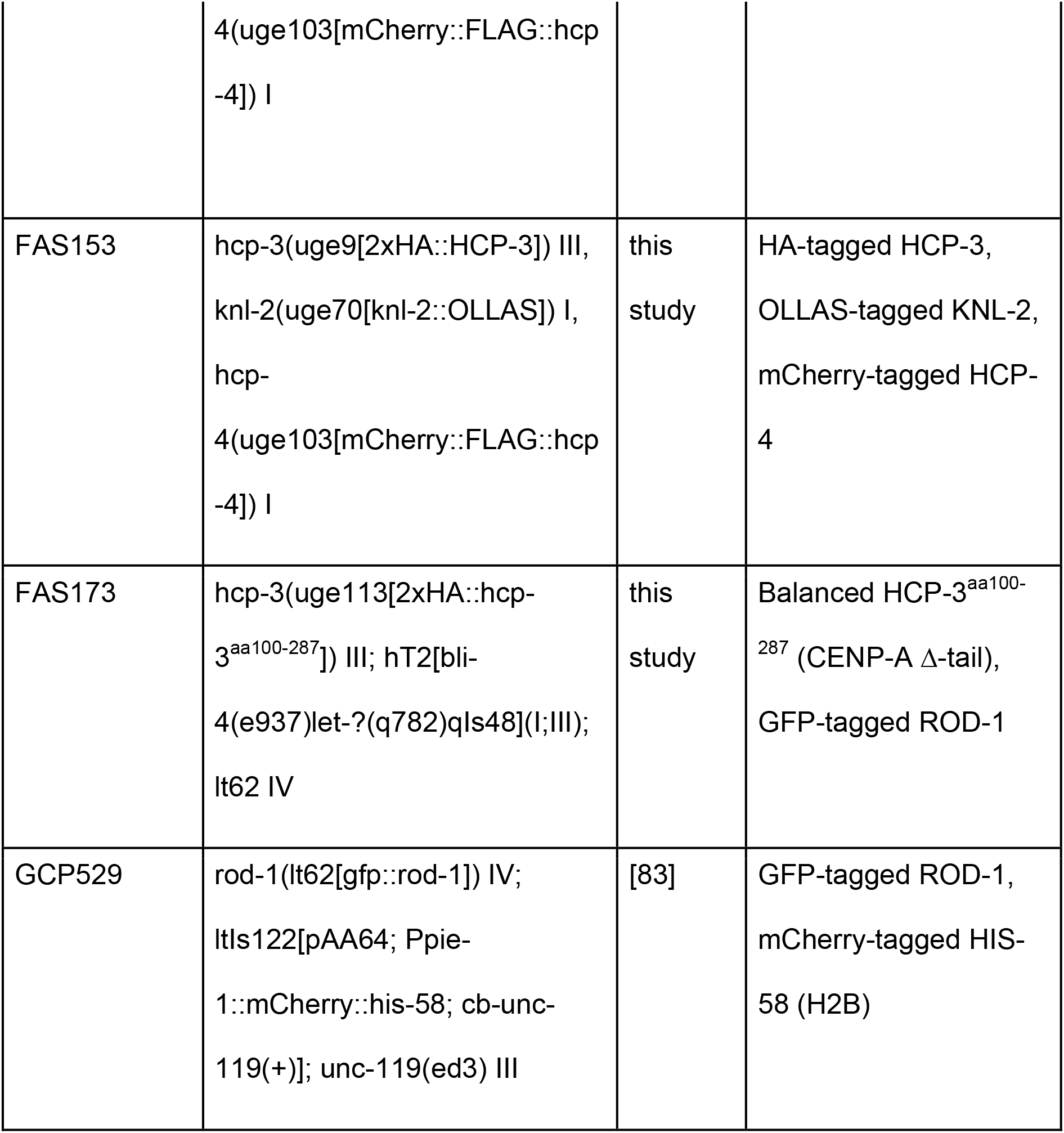
*C. elegans* strains generated or used in this study.

## References

1. Fukagawa T, Earnshaw WC. The centromere: chromatin foundation for the kinetochore machinery. Dev Cell. 2014;30: 496–508.

2. McKinley KL, Cheeseman IM. The molecular basis for centromere identity and function. Nat Rev Mol Cell Biol. 2016;17: 16–29.

3. Dunleavy EM, Almouzni G, Karpen GH. H3.3 is deposited at centromeres in S phase as a placeholder for newly assembled CENP-A in G1phase. Nucleus. 2011. pp. 146–157. doi:10.4161/nucl.2.2.15211

4. Malik HS, Henikoff S. Phylogenomics of the nucleosome. Nat Struct Biol. 2003;10: 882–891.

5. Talbert PB, Bryson TD, Henikoff S. Adaptive evolution of centromere proteins in plants and animals. J Biol. 2004;3: 18.

6. Schueler MG, Swanson W, Thomas PJ, NISC Comparative Sequencing Program, Green ED. Adaptive evolution of foundation kinetochore proteins in primates. Mol Biol Evol. 2010;27: 1585–1597.

7. Mariño-Ramírez L, Kann MG, Shoemaker BA, Landsman D. Histone structure and nucleosome stability. Expert Rev Proteomics. 2005;2: 719–729.

8. Chen C-C, Mellone BG. Chromatin assembly: Journey to the CENter of the chromosome. J Cell Biol. 2016;214: 13–24.

9. Stellfox ME, Bailey AO, Foltz DR. Putting CENP-A in its place. Cell Mol Life Sci. 2013;70: 387–406.

10. Jansen LET, Black BE, Foltz DR, Cleveland DW. Propagation of centromeric chromatin requires exit from mitosis. The Journal of Cell Biology. 2007. pp. 795–805. doi:10.1083/jcb.200701066

11. Hayashi T, Fujita Y, Iwasaki O, Adachi Y, Takahashi K, Yanagida M. Mis16 and Mis18 are required for CENP-A loading and histone deacetylation at centromeres. Cell. 2004;118: 715–729.

12. Fujita Y, Hayashi T, Kiyomitsu T, Toyoda Y, Kokubu A, Obuse C, et al. Priming of centromere for CENP-A recruitment by human hMis18alpha, hMis18beta, and M18BP1. Dev Cell. 2007;12: 17–30.

13. Kwon M-S, Hori T, Okada M, Fukagawa T. CENP-C is involved in chromosome segregation, mitotic checkpoint function, and kinetochore assembly. Mol Biol Cell. 2007;18: 2155–2168.

14. Milks KJ, Moree B, Straight AF. Dissection of CENP-C–directed Centromere and Kinetochore Assembly. Molecular Biology of the Cell. 2009. pp. 4246–4255. doi:10.1091/mbc.e09-05-0378

15. Foltz DR, Jansen LET, Bailey AO, Yates JR 3rd, Bassett EA, Wood S, et al. Centromere-specific assembly of CENP-a nucleosomes is mediated by HJURP. Cell. 2009;137: 472–484.

16. Dunleavy EM, Roche D, Tagami H, Lacoste N, Ray-Gallet D, Nakamura Y, et al. HJURP is a cell-cycle-dependent maintenance and deposition factor of CENP-A at centromeres. Cell. 2009;137: 485–497.

17. Chen C-C, Dechassa ML, Bettini E, Ledoux MB, Belisario C, Heun P, et al. CAL1 is the Drosophila CENP-A assembly factor. J Cell Biol. 2014;204: 313–329.

18. Shukla M, Tong P, White SA, Singh PP, Reid AM, Catania S, et al. Centromere DNA Destabilizes H3 Nucleosomes to Promote CENP-A Deposition during the Cell Cycle. Curr Biol. 2018;28: 3924–3936.e4.

19. Choi ES, Strålfors A, Catania S, Castillo AG, Svensson JP, Pidoux AL, et al. Factors that promote H3 chromatin integrity during transcription prevent promiscuous deposition of CENP-A(Cnp1) in fission yeast. PLoS Genet. 2012;8: e1002985.

20. Chen C-C, Bowers S, Lipinszki Z, Palladino J, Trusiak S, Bettini E, et al. Establishment of Centromeric Chromatin by the CENP-A Assembly Factor CAL1 Requires FACT-Mediated Transcription. Dev Cell. 2015;34: 73–84.

21. Bobkov GOM, Gilbert N, Heun P. Centromere transcription allows CENP-A to transit from chromatin association to stable incorporation. J Cell Biol. 2018;217: 1957–1972.

22. Prosée RF, Wenda JM, Steiner FA. Adaptations for centromere function in meiosis. Essays Biochem. 2020. doi:10.1042/EBC20190076

23. Raychaudhuri N, Dubruille R, Orsi GA, Bagheri HC, Loppin B, Lehner CF. Transgenerational propagation and quantitative maintenance of paternal centromeres depends on Cid/Cenp-A presence in Drosophila sperm. PLoS Biol. 2012;10: e1001434.

24. Swartz SZ, McKay LS, Su K-C, Bury L, Padeganeh A, Maddox PS, et al. Quiescent Cells Actively Replenish CENP-A Nucleosomes to Maintain Centromere Identity and Proliferative Potential. Dev Cell. 2019;51: 35–48.e7.

25. Smoak EM, Stein P, Schultz RM, Lampson MA, Black BE. Long-Term Retention of CENP-A Nucleosomes in Mammalian Oocytes Underpins Transgenerational Inheritance of Centromere Identity. Curr Biol. 2016;26: 1110–1116.

26. Gassmann R, Rechtsteiner A, Yuen KW, Muroyama A, Egelhofer T, Gaydos L, et al. An inverse relationship to germline transcription defines centromeric chromatin in C. elegans. Nature. 2012;484: 534–537.

27. Ingouff M, Rademacher S, Holec S, Soljić L, Xin N, Readshaw A, et al. Zygotic resetting of the HISTONE 3 variant repertoire participates in epigenetic reprogramming in Arabidopsis. Curr Biol. 2010;20: 2137–2143.

28. Monen J, Maddox PS, Hyndman F, Oegema K, Desai A. Differential role of CENP-A in the segregation of holocentric C. elegans chromosomes during meiosis and mitosis. Nat Cell Biol. 2005;7: 1248–1255.

29. Steiner FA, Henikoff S. Diversity in the organization of centromeric chromatin. Curr Opin Genet Dev. 2015;31: 28–35.

30. Maddox PS, Hyndman F, Monen J, Oegema K, Desai A. Functional genomics identifies a Myb domain-containing protein family required for assembly of CENP-A chromatin. J Cell Biol. 2007;176: 757–763.

31. Lee BCH, Lin Z, Yuen KWY. RbAp46/48LIN-53 Is Required for Holocentromere Assembly in Caenorhabditis elegans. Cell Reports. 2016. pp. 1819–1828. doi:10.1016/j.celrep.2016.01.065

32. Oegema K, Desai A, Rybina S, Kirkham M, Hyman AA. Functional analysis of kinetochore assembly in Caenorhabditis elegans. J Cell Biol. 2001;153: 1209–1226.

33. Boxem M, Maliga Z, Klitgord N, Li N, Lemmens I, Mana M, et al. A protein domain-based interactome network for C. elegans early embryogenesis. Cell. 2008;134: 534–545.

34. Delaney K, Mailler J, Wenda JM, Gabus C, Steiner FA. Differential Expression of Histone H3.3 Genes and Their Role in Modulating Temperature Stress Response in. Genetics. 2018;209: 551–565.

35. Ooi SL, Priess JR, Henikoff S. Histone H3.3 variant dynamics in the germline of Caenorhabditis elegans. PLoS Genet. 2006;2: e97.

36. Buchwitz BJ, Ahmad K, Moore LL, Roth MB, Henikoff S. A histone-H3-like protein in C. elegans. Nature. 1999. pp. 547–548. doi:10.1038/44062

37. Zhang L, Ward JD, Cheng Z, Dernburg AF. The auxin-inducible degradation (AID) system enables versatile conditional protein depletion in C. elegans. Development. 2015;142: 4374–4384.

38. Bodor DL, Valente LP, Mata JF, Black BE, Jansen LET. Assembly in G1 phase and long-term stability are unique intrinsic features of CENP-A nucleosomes. Mol Biol Cell. 2013;24: 923–932.

39. Mellone BG, Grive KJ, Shteyn V, Bowers SR, Oderberg I, Karpen GH. Assembly of Drosophila centromeric chromatin proteins during mitosis. PLoS Genet. 2011;7: e1002068.

40. Dunleavy EM, Beier NL, Gorgescu W, Tang J, Costes SV, Karpen GH. The cell cycle timing of centromeric chromatin assembly in Drosophila meiosis is distinct from mitosis yet requires CAL1 and CENP-C. PLoS Biol. 2012;10: e1001460.

41. Barnhart MC, Kuich PHJL, Stellfox ME, Ward JA, Bassett EA, Black BE, et al. HJURP is a CENP-A chromatin assembly factor sufficient to form a functional de novo kinetochore. J Cell Biol. 2011;194: 229–243.

42. Gascoigne KE, Takeuchi K, Suzuki A, Hori T, Fukagawa T, Cheeseman IM. Induced ectopic kinetochore assembly bypasses the requirement for CENP-A nucleosomes. Cell. 2011;145: 410–422.

43. Hori T, Shang W-H, Takeuchi K, Fukagawa T. The CCAN recruits CENP-A to the centromere and forms the structural core for kinetochore assembly. J Cell Biol. 2013;200: 45–60.

44. Moree B, Meyer CB, Fuller CJ, Straight AF. CENP-C recruits M18BP1 to centromeres to promote CENP-A chromatin assembly. J Cell Biol. 2011;194: 855–871.

45. Hori T, Shang W-H, Hara M, Ariyoshi M, Arimura Y, Fujita R, et al. Association of M18BP1/KNL2 with CENP-A Nucleosome Is Essential for Centromere Formation in Non-mammalian Vertebrates. Dev Cell. 2017;42: 181–189.e3.

46. French BT, Westhorpe FG, Limouse C, Straight AF. Xenopus laevis M18BP1 Directly Binds Existing CENP-A Nucleosomes to Promote Centromeric Chromatin Assembly. Dev Cell. 2017;42: 190–199.e10.

47. Lermontova I, Kuhlmann M, Friedel S, Rutten T, Heckmann S, Sandmann M, et al. Arabidopsis kinetochore null2 is an upstream component for centromeric histone H3 variant cenH3 deposition at centromeres. Plant Cell. 2013;25: 3389–3404.

48. Kral L. Possible identification of CENP-C in fish and the presence of the CENP-C motif in M18BP1 of vertebrates. F1000Res. 2015;4: 474.

49. Sandmann M, Talbert P, Demidov D, Kuhlmann M, Rutten T, Conrad U, et al. Targeting of Arabidopsis KNL2 to Centromeres Depends on the Conserved CENPC-k Motif in Its C Terminus. Plant Cell. 2017;29: 144–155.

50. Mach J. A Tale of Two CENPCs: Centromere Localization of KINETOCHORE NULL2 and CENP-C. The Plant cell. 2017. pp. 2–3.

51. Fachinetti D, Diego Folco H, Nechemia-Arbely Y, Valente LP, Nguyen K, Wong AJ, et al. A two-step mechanism for epigenetic specification of centromere identity and function. Nature Cell Biology. 2013. pp. 1056–1066. doi:10.1038/ncb2805

52. Logsdon GA, Barrey EJ, Bassett EA, DeNizio JE, Guo LY, Panchenko T, et al. Both tails and the centromere targeting domain of CENP-A are required for centromere establishment. J Cell Biol. 2015;208: 521–531.

53. Bailey AO, Panchenko T, Sathyan KM, Petkowski JJ, Pai P-J, Bai DL, et al. Posttranslational modification of CENP-A influences the conformation of centromeric chromatin. Proc Natl Acad Sci U S A. 2013;110: 11827–11832.

54. Sathyan KM, Fachinetti D, Foltz DR. α-amino trimethylation of CENP-A by NRMT is required for full recruitment of the centromere. Nat Commun. 2017;8: 14678.

55. Barra V, Logsdon GA, Scelfo A, Hoffmann S, Hervé S, Aslanian A, et al. Phosphorylation of CENP-A on serine 7 does not control centromere function. Nat Commun. 2019;10: 175.

56. Tachiwana H, Müller S, Blümer J, Klare K, Musacchio A, Almouzni G. HJURP involvement in de novo CenH3(CENP-A) and CENP-C recruitment. Cell Rep. 2015;11: 22–32.

57. Folco HD, Campbell CS, May KM, Espinoza CA, Oegema K, Hardwick KG, et al. The CENP-A N-tail confers epigenetic stability to centromeres via the CENP-T branch of the CCAN in fission yeast. Curr Biol. 2015;25: 348–356.

58. Takada M, Zhang W, Suzuki A, Kuroda TS, Yu Z, Inuzuka H, et al. FBW7 Loss Promotes Chromosomal Instability and Tumorigenesis via Cyclin E1/CDK2-Mediated Phosphorylation of CENP-A. Cancer Res. 2017;77: 4881–4893.

59. Zeitlin SG, Shelby RD, Sullivan KF. CENP-A is phosphorylated by Aurora B kinase and plays an unexpected role in completion of cytokinesis. J Cell Biol. 2001;155: 1147–1157.

60. Kunitoku N, Sasayama T, Marumoto T, Zhang D, Honda S, Kobayashi O, et al. CENP-A Phosphorylation by Aurora-A in Prophase Is Required for Enrichment of Aurora-B at Inner Centromeres and for Kinetochore Function. Developmental Cell. 2003. pp. 853–864. doi:10.1016/s1534-5807(03)00364-2

61. Goutte-Gattat D, Shuaib M, Ouararhni K, Gautier T, Skoufias DA, Hamiche A, et al. Phosphorylation of the CENP-A amino-terminus in mitotic centromeric chromatin is required for kinetochore function. Proc Natl Acad Sci U S A. 2013;110: 8579–8584.

62. Huang A, Kremser L, Schuler F, Wilflingseder D, Lindner H, Geley S, et al. Phosphorylation of Drosophila CENP-A on serine 20 regulates protein turn-over and centromere-specific loading. Nucleic Acids Res. 2019;47: 10754–10770.

63. Srivastava S, Foltz DR. Posttranslational modifications of CENP-A: marks of distinction. Chromosoma. 2018. pp. 279–290. doi:10.1007/s00412-018-0665-x

64. Lermontova I, Koroleva O, Rutten T, Fuchs J, Schubert V, Moraes I, et al. Knockdown of CENH3 in Arabidopsis reduces mitotic divisions and causes sterility by disturbed meiotic chromosome segregation. The Plant Journal. 2011. pp. 40–50. doi:10.1111/j.1365-313x.2011.04664.x

65. Ravi M, Shibata F, Ramahi JS, Nagaki K, Chen C, Murata M, et al. Meiosis-Specific Loading of the Centromere-Specific Histone CENH3 in Arabidopsis thaliana. PLoS Genetics. 2011. p. e1002121. doi:10.1371/journal.pgen.1002121

66. Desai A, Rybina S, Müller-Reichert T, Shevchenko A, Shevchenko A, Hyman A, Oegema K. KNL-1 directs assembly of the microtubule-binding interface of the kinetochore in C. elegans. Genes & Development. 2003. pp. 2421–2435. doi:10.1101/gad.1126303

67. Cheeseman IM, Niessen S, Anderson S, Hyndman F, Yates JR 3rd, Oegema K, et al. A conserved protein network controls assembly of the outer kinetochore and its ability to sustain tension. Genes Dev. 2004;18: 2255–2268.

68. Luschnig S, Moussian B, Krauss J, Desjeux I, Perkovic J, Nüsslein-Volhard C. An F1 genetic screen for maternal-effect mutations affecting embryonic pattern formation in Drosophila melanogaster. Genetics. 2004;167: 325–342.

69. Hekimi S, Boutis P, Lakowski B. Viable maternal-effect mutations that affect the development of the nematode Caenorhabditis elegans. Genetics. 1995;141: 1351–1364.

70. Mango SE, Thorpe CJ, Martin PR, Chamberlain SH, Bowerman B. Two maternal genes, apx-1 and pie-1, are required to distinguish the fates of equivalent blastomeres in the early Caenorhabditis elegans embryo. Development. 1994;120: 2305–2315.

71. Mello CC, Draper BW, Prless JR. The maternal genes apx-1 and glp-1 and establishment of dorsal-ventral polarity in the early C. elegans embryo. Cell. 1994. pp. 95–106. doi:10.1016/0092-8674(94)90238-0

72. Bénard C, McCright B, Zhang Y, Felkai S, Lakowski B, Hekimi S. The C. elegans maternal-effect gene clk-2 is essential for embryonic development, encodes a protein homologous to yeast Tel2p and affects telomere length. Development. 2001;128: 4045–4055.

73. Mains PE, Sulston IA, Wood WB. Dominant maternal-effect mutations causing embryonic lethality in Caenorhabditis elegans. Genetics. 1990;125: 351–369.

74. Cenik ES, Meng X, Tang NH, Hall RN, Arribere JA, Cenik C, et al. Maternal Ribosomes Are Sufficient for Tissue Diversification during Embryonic Development in C. elegans. Dev Cell. 2019;48: 811–826.e6.

75. Woodhouse RM, Ashe A. How do histone modifications contribute to transgenerational epigenetic inheritance in C. elegans? Biochemical Society Transactions. 2020. pp. 1019–1034. doi:10.1042/bst20190944

76. Gaydos LJ, Wang W, Strome S. Gene repression. H3K27me and PRC2 transmit a memory of repression across generations and during development. Science. 2014;345: 1515–1518.

77. Kaneshiro KR, Rechtsteiner A, Strome S. Sperm-inherited H3K27me3 impacts offspring transcription and development in C. elegans. Nature Communications. 2019. doi:10.1038/s41467-019-09141-w

78. Burkhart KB, Sando SR, Corrionero A, Horvitz HR. H3.3 Nucleosome Assembly Mutants Display a Late-Onset Maternal Effect. Curr Biol. 2020;30: 2343–2352.e3.

79. Arribere JA, Bell RT, Fu BXH, Artiles KL, Hartman PS, Fire AZ. Efficient marker-free recovery of custom genetic modifications with CRISPR/Cas9 in Caenorhabditis elegans. Genetics. 2014;198: 837–846.

80. Kamath RS, Fraser AG, Dong Y, Poulin G, Durbin R, Gotta M, et al. Systematic functional analysis of the Caenorhabditis elegans genome using RNAi. Nature. 2003;421: 231–237.

81. Schindelin J, Arganda-Carreras I, Frise E, Kaynig V, Longair M, Pietzsch T, et al. Fiji: an open-source platform for biological-image analysis. Nat Methods. 2012;9: 676–682.

82. Kress E, Schwager F, Holtackers R, Seiler J, Prodon F, Zanin E, et al. The UBXN-2/p37/p47 adaptors of CDC-48/p97 regulate mitosis by limiting the centrosomal recruitment of Aurora A. J Cell Biol. 2013;201: 559–575.

83. Pereira C, Reis RM, Gama JB, Celestino R, Cheerambathur DK, Carvalho AX, et al. Self-Assembly of the RZZ Complex into Filaments Drives Kinetochore Expansion in the Absence of Microtubule Attachment. Curr Biol. 2018;28: 3408–3421.e8.

